# Traits and hidden states: is self-fertilization associated with rates of diversification across mating and sexual systems?

**DOI:** 10.64898/2026.06.18.733044

**Authors:** Elena M. Meyer, Michael S. Rosenberg, Bret M. Boyd, Andrew J. Eckert

## Abstract

Angiosperms, the most diverse group of land plants, possess considerable variation in sexual expression and mating strategy. Reproductive biology is a key determinant of fitness, and may help answer a persistent question in evolutionary biology: why are some areas of the tree of life more species-rich than others? State-dependent speciation-extinction methods (SSEs), which associate traits with patterns of diversification, can be used here. Previously, SSEs have been used within families to investigate the hypothesis that selfing is an evolutionary “dead end.” To the best of the authors’ knowledge, no study has looked across families, which would increase power to more generally test this hypothesis. Here, we examine the impact of 1) mating system and 2) sexual system on diversification across 18 phylogenetically diverse families. We also discuss how more recent advances in SSE models (i.e., “hidden state” models) influence our interpretation of these patterns and evaluate how the relationship between mating and sexual systems can be leveraged to gain insight into the impact of reproductive biology on evolutionary outcomes. We find that the mating system does not better explain patterns of diversification when compared to null models, but the sexual system often does. We also find family-level heterogeneity in our results, which suggests conclusions drawn from studies on individual families may not be consistent with any broader trend.

## Introduction

Understanding how angiosperms were able to rapidly diversify from a point of relative rarity in the early and mid-Cretaceous period to become today’s dominant form of plant life has been a guiding question in botany and evolutionary biology since Darwin (Friedman 2009). Across all parts of the tree of life, expansion or contraction of clades is an ongoing and fluid process shaped by speciation and extinction (Darwin 1859; Raup 1995, Kopperud et al. 2026). (Stankowski and Ravinet 2021). Examining how shared traits across species impact evolutionary outcomes can help explain the underlying forces that have shaped the tree of life. Thus, the comparative method (i.e., using comparison between species to draw generalizable conclusions; reviewed Harvey and Pagel 1991), when used in a phylogenetically informed manner (Felsenstein 1985), can offer profound insights into how and why diversity varies between clades. Prior studies have shown that diversification rates vary across the tree of life and that variation is often based on differing organismal traits (Mitter et al., 1986; Scholl and Wiens, 2016). Uncovering patterns in how specific traits impact diversification is a key aim of macroevolutionary studies (Maddison et al. 2007; Jablonski 2017). Toward this aim, phylogenetic comparative methods, such as state-dependent speciation-extinction (SSE) models, offer a vital way to examine these dynamics on deep timescales by linking trait evolution with changes in diversification rates (Maddison et al. 2007, reviewed by Harmon 2018).

Reproduction is fundamental to fitness (Darwin 1859, Fisher 1930), and different reproductive systems could influence speciation and extinction rates. Plants are an excellent system in which to test this idea. Plants reproduce with a variety of mechanisms (Barrett 2002), and angiosperms, more specifically, use different partitioning of male and female gamete-producing structures between or within flowers, which also affect realized mating systems (i.e., dioecious species are definitionally outcrossing). Understanding the influence of reproductive characters on diversification is key to understanding angiosperm diversity (Darwin 1876, compiled 1903; Friedman 2009, Sauquet and Magallón 2018). Unsurprisingly, the variation in floral form and function associated with mating was of interest to Darwin (Darwin 1862, 1876, 1877; reviewed in Barrett 2010). Despite a long history of scientific investigation, the study of plant reproductive biology continues to yield new evolutionary insights.

Plant reproductive biology includes variation in both mating system (i.e., how mating takes place within the individual or between individuals in a population) and sexual system (i.e., how sexual characteristics vary between individuals). In plant mating systems, some plants self-fertilize, while others only reproduce by outcrossing. Some plants, however, display mixed mating, meaning that they are capable of both outcrossing and self-fertilization. Approximately 58% of angiosperms with known mating systems are self-compatible (Meyer et al. 2025), although this data is known to be far from random (Goodwillie et al. 2005), and likely over- estimates the true frequency of selfing in angiosperms (Igić and Kohn 2006, Meyer et al. 2025).

In angiosperms, sexual system can differ based on floral characteristics. Individuals can be hermaphrodites, possessing both male and female organs on a single flower. Alternatively, some species exhibit a separation of the male and female parts among flowers on a single individual (monoecy) or male and female individuals (dioecy). Based on the large-scale review in Wang et al. (2021), we can approximate hermaphroditism to be present in 76% of plants with known sexual systems, with monoecy present in approximately 16%, and dioecy in 8%. While mating systems and sexual systems are often treated separately in the literature (e.g., Goodwillie et al. 2005), both traits impact reproductive dynamics. For example, individuals that are hermaphroditic have the fewest physiological barriers to self-fertilization, as the male and female parts of the flower are not separated. While selfing within the individual is possible for monoecious individuals, it would require the facilitation of pollen transfer from a male to female flower, presenting a potential barrier. Finally, for dioecious species, selfing is physiologically impossible due to the separation of the sexes across individuals. Thus, both direct information on selfing from mating system studies and indirect information from knowledge of sexual systems based on morphology can be used to elucidate the effect of self-fertilization in angiosperm evolution (see also: Meyer et al. 2025). Both systems can be leveraged for insights into how plant species evolve.

Studies across the breadth of angiosperm diversity suggest selfing leads to increased rates of extinction and thus lower diversification rates (Ferrer and Good 2012; Wright et al. 2013). SSEs provide a nuanced picture of evolutionary dynamics by joint estimation of speciation, extinction, and rates of transition (Maddison et al., 2007). SSEs have been applied to some individual angiosperm families to examine mating system dynamics, yielding mixed evidence that self-fertilization is associated with lower diversification rates (Solanaceae: Goldberg et al. 2010; Primulaceae: de Vos et al. 2014; Onagraceae: Freyman and Höhna 2019). However, no single study has included multiple angiosperm families utilizing SSEs. This approach is vital to uncover – or dispute – a generalizable macroevolutionary pattern of self-fertilization as a driver of diversification rates.

Self-fertilization has been of particular interest to evolutionary biologists, and has been hypothesized to have a negative impact on diversification rates (i.e., “dead end” or “blind alley” hypothesis; Stebbins 1957). However, only a limited number of studies have examined the impact of angiosperm sexual systems on diversification rates (discussed by Sabath et al. 2016). In addition, mating and sexual systems have never been looked at in conjunction with each other. Since data about reproductive traits in angiosperms is limited (Meyer et al. 2025), this integrative approach allows for us to address a broader number of species and fosters a more complete understanding of overall reproductive dynamics, especially since sexual system is at least partially informative about mating system.

To illuminate how reproductive traits influence diversification rates, we utilize SSE methods, which link speciation and extinction—and thus net diversification rates—with changes in trait values (Maddison et al. 2007). The simplest of these methods, BiSSE (Binary State Speciation and Extinction; Maddison et al. 2007) uses binary traits, which is suitable for working with designations of either self-incompatibility or self-compatibility. Since the initial publication of BiSSE, a variety of expansions to the method to allow for non-binary data have proliferated (reviewed in Marie-Orleach et al. 2024). One of these methods, MuSSE (Multistate Speciation and Extinction; FitzJohn, 2012), allows for more than two states, such as is present with sexual systems. This expanded framework allows for examination of both binary multi-state characters, such as mating system and sexual system data.

One major area of recent research in SSE models is the generation of appropriately complex null models. If null models are overly trivial, rejecting these null models is not a strong indication that the focal trait meaningfully impacts diversification rates. For example, in a typical null model for BiSSE, the focal trait does not impact speciation-extinction patterns, thus requiring equal rates of diversification (Revell and Harmon 2022). However, even if the trait of interest does not affect diversification rates, a model incorporating rate variation is often a better fit for realistically complex datasets (Beaulieu and O’Meara 2016). This is especially important since net diversification rates are known to vary across groups of plants (Scholl and Wiens 2016) and the entire tree of life (Kopperud et al. 2026). In other words, if there is variation in net diversification rates in a phylogeny, regardless of their covariation with the traits of interest, it is often easy to reject a null model which assumes rate constancy (Rabosky and Goldberg 2015, Beaulieu and O’Meara 2016, Caetano et al. 2018).

One potential solution to this problem is the HiSSE framework as proposed in Beaulieu and O’Meara (2016). HiSSE introduces “hidden states” in addition to observed states, which would allow for differing diversification rates, and the generation of appropriately complex models. The general structure of a HiSSE model is reviewed in **Figure 1**.

**Figure 1.**
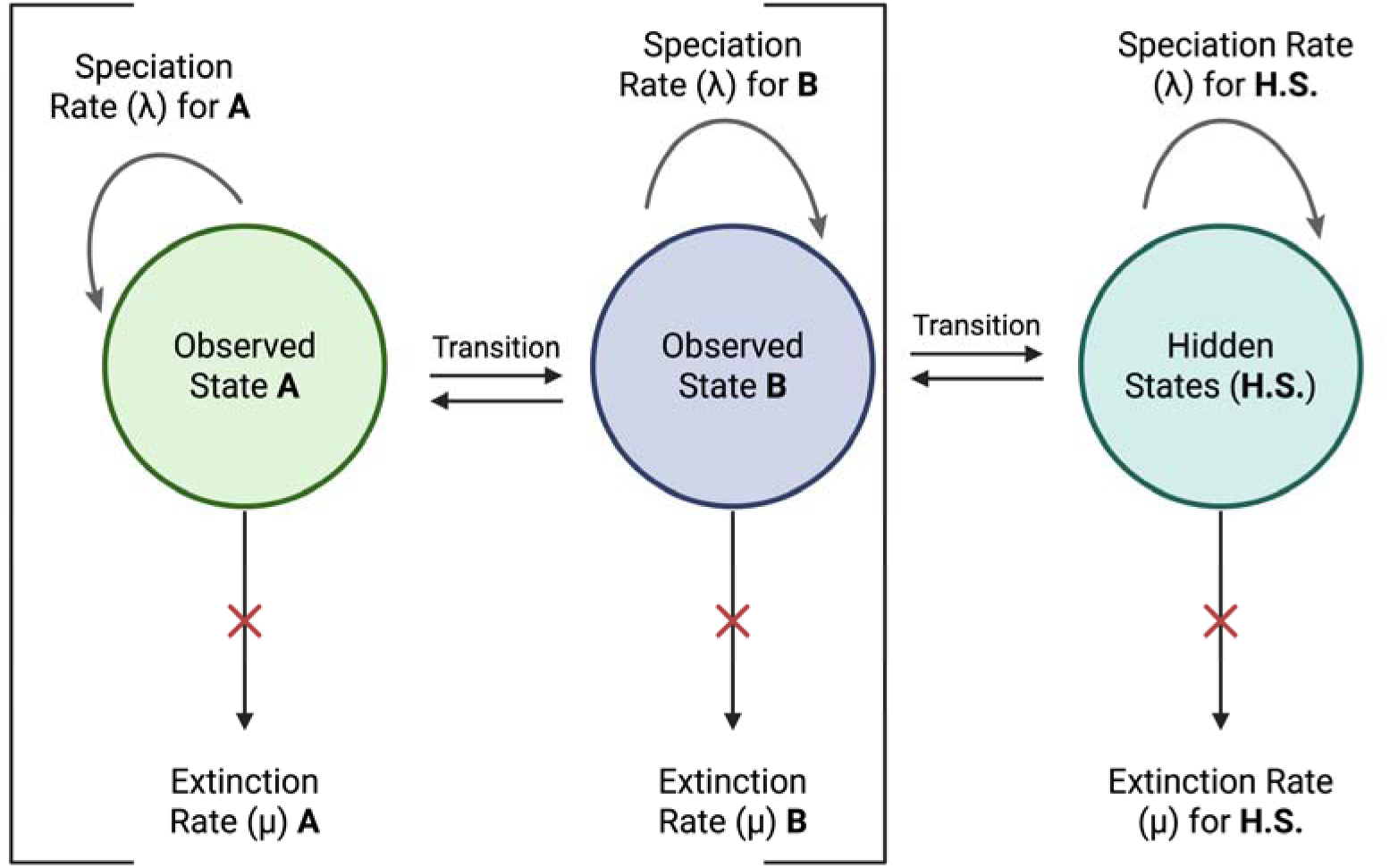
Diagram showing a simple example of how state-dependent speciation-extinction models work. Here, a two-state model is shown with transition rates between State A and State B allowed to vary freely with each other. States A and B are also allowed to vary freely with one or more hidden states. Rates of speciation and extinction are also included for each state and for the hidden state(s). Note here that the hidden state(s) can transition between both State A and B (as indicated by the brackets). Figure created with Biorender.

To address the problem of rejecting an overly trivial null, Beaulieu and O’Meara (2016) proposed a set of character-independent (CID) models which can be used as complex null models for BiSSE-like and HiSSE-like models. These models are a character-independent, two-rate model with two state levels (CID-2) and a character-independent four-rate model with a four-level state (CID-4). Note here that these numbers correspond to a two-state model; in expansions of the method where multi-state HiSSE was used (i.e., Nakov et al. 2018), the appropriate null for a full HiSSE model would have additional free parameters to match the number of measured states. These advancements should add nuance to the overall picture relative to previous studies based on SSEs.

Here, we utilize a broadly distributed set of families (*n* = 18) sampled from across the angiosperm phylogeny to characterize the impact of mating systems and sexual systems on rates of speciation, extinction, and diversification over time. We use this approach to investigate two key questions: 1) do reproductive traits affect rates of net diversification across families, and 2) is this impact consistent across families? We then discuss the implications of our findings for our understanding of the evolution of selfing and associated reproductive characters, and how the advent of hidden-state phylogenetic models has led to new results changing and deepening our understanding of how mating systems evolve.

## Methods

### Data sourcing and selection

All analyses for this paper were conducted using R version 4.3.0 (R Core Team, 2023). We sourced our phylogenetic information from a recent angiosperm-wide mega-phylogeny which included 79,881 species (Smith and Brown 2018, the GBOTB tree). To populate tips on the phylogeny with traits, we utilized binary-level mating system data (either selfing or outcrossing) from Meyer et al. (2025), and three-state sexual system data from Wang et al. (2021). All names were standardized to World Flora Online using the package `U.Taxonstrand` version 1.1.3 (Zhang and Qian 2023).

We filtered our trait data for tree tips by family, and selected families which had 1) a minimum of 90 matches between species represented on the tree and trait data, and 2) variation in the traits of interest. Since our primary goal was to test for speciation rate asymmetry associated with reproductive system traits, this modest minimum sample size was deemed likely to still have sufficient statistical power for our specific goals (Gamisch 2016). Families which either had limited representation in our data or no appropriate trait-level variation in the focal traits were excluded. The resultant families, as well as the number of tips in each state, are listed in **Table 1**. We then pruned the larger phylogeny to obtain rooted trees for our 18 sampled families by dropping tips for species without trait data for each family using the `ape` package ver. 5.8 (Paradis and Schliep 2019). The final product was a set of 20 trimmed trees with complete trait data populating the tips, corresponding to three mating system-matched trees and 17 sexual-system matched trees (Solanaceae and Asteraceae generated trait-matched trees under our criteria for both traits, while Brassicaceae matched sexual system but not mating system).

**Table 1.** A summary of the scope of this paper: the column **Family** indicates the taxonomic family studied; **Type** indicates if the trait was binary state or multistate, **Tips** indicates the number of matched tips on the phylogeny with data; **Size** is the approximate number of species in the family (as given in Christenhusz and Byng 2016); **% Cover** is the proportion of the family for which there is data in our sample. The remaining columns indicate the number of tips for each trait state: dioecious, monoecious, hermaphroditic, self-compatible, and self-incompatible, respectively.

| **Family** | **Type** | **Tips** | **Size** | **% Cover** | **Dio. tips** | **Herm.** | **Mono.** | **SC** | **SI** |
| --- | --- | --- | --- | --- | --- | --- | --- | --- | --- |
| Apiaceae | Multi | 375 | 3575 | 10.49 | 23 | 301 | 51 | - | - |
| Araceae | Multi | 221 | 3750 | 5.89 | 18 | 54 | 149 | - | - |
| Asteraceae | Binary | 477 | 24700 | 1.93 | - | - | - | 183 | 294 |
| Asteraceae | Multi | 2214 | 24700 | 8.96 | 79 | 909 | 1229 | - | - |
| Brassicaceae | Binary | 200 | 3628 | 5.51 | - | - | - | 137 | 63 |
| Cucurbitaceae | Binary | 215 | 965 | 22.28 | 125 | 0 | 90 | - | - |
| Euphorbiaceae | Multi | 836 | 6252 | 13.37 | 70 | 3 | 763 | - | - |
| Lamiaceae | Binary | 489 | 7530 | 6.49 | 25 | 464 | 0 | - | - |
| Lauraceae | Multi | 214 | 2850 | 7.51 | 97 | 116 | 1 | - | - |
| Loranthaceae | Multi | 97 | 1050 | 9.24 | 5 | 91 | 1 | - | - |
| Moraceae | Binary | 232 | 1180 | 19.66 | 91 | 0 | 141 | - | - |
| Piperaceae | Multi | 105 | 3700 | 2.84 | 23 | 80 | 2 | - | - |
| Poaceae | Multi | 866 | 12000 | 7.22 | 6 | 655 | 205 | - | - |
| Rosaceae | Multi | 641 | 2950 | 21.73 | 12 | 627 | 2 | - | - |
| Rutaceae | Multi | 90 | 2070 | 4.35 | 50 | 33 | 7 | - | - |
| Salicaceae | Multi | 154 | 1220 | 12.62 | 145 | 8 | 1 | - | - |
| Solanaceae | Binary | 285 | 2600 | 10.96 | - | - | - | 193 | 92 |
| Solanaceae | Multi | 471 | 2600 | 18.12 | 5 | 453 | 13 | - | - |
| Urticaceae | Multi | 111 | 2625 | 4.23 | 40 | 2 | 69 | - | - |
| Vitaceae | Binary | 90 | 910 | 9.89 | 64 | 26 | 0 | - | - |

### Analysis of association between binary mating system and diversification

We used the `hisse` package version 2.1.11 (Beaulieu and O’Meara 2016) to jointly estimate rates of speciation and extinction conditioned on the traits and the tree provided for the binary mating system data. Note that HiSSE calculates extinction fraction and species turnover, rather than directly calculating speciation and extinction rates like prior SSE models based on BiSSE (Maddison et al. 2007). However, speciation and extinction rates can easily be back-calculated from these values.

For our binary-state mating system data, we implemented two types of models: 1) character independent diversification models (CID), and 2) character-dependent SSE models with 0-4 correlated hidden states (i.e., BiSSE and HiSSE).

CID models act as null models for state-dependent diversification and can be scaled to match complexity with BiSSE and HiSSE models (Beaulieu and O’Meara, 2016, Revell and Harmon 2022). While independent of the focal traits (i.e., characters), as the name suggests, these models allow for rate variation in speciation and extinction rates associated with the hidden states This is vital to avoid pitfalls associated with early iterations of the BiSSE framework where any form of rate variation could lead to rejecting the null model (see Rabosky and Goldberg 2015 and Beaulieu and O’Meara 2016). We broadly followed Revell and Harmon (2022) for guidance in fixing parameters (e.g., transition rates between states). To calculate sampling fraction estimates for character states, we used the complete datasets from Wang et al. (2021) and Meyer et al. (2025), and family size estimates from Christenhusz and Byng (2016) to estimate the family-level prevalence of each character state and compare against our tree- matched datasets.

Our character-independent models were comprised of three types:

1. A minimal null model, CID, in which each of the estimated rates of speciation and extinction were held equal for selfing and outcrossing, yielding a single rate estimation for speciation, extinction, forward transitions, and back transitions (*k* = 4). In other words, a constant birth-death model or the “normal” BiSSE null model (Maddison et al. 2007; Revell and Harmon 2022).
2. A character-independent null model, CID-2, which is otherwise similar in complexity to the character-dependent BiSSE model with two characters, both with equal rates for speciation, extinction, and a single rate for forward and backward transitions (*k* = 5).
3. A four-state character-independent model, CID-4, which is otherwise similar to a “full” HiSSE model; this model has rates associated with speciation and extinction parameters for four characters (yielding eight estimates of these parameters) plus a single rate for transitions (*k* = 9).

Character-dependent models incorporate character trait data to estimate diversification parameters. Our character-dependent models were of the following two types:

1. A two-state character dependent model with no hidden states, which is equivalent to a standard BiSSE model: speciation and extinction rates for two binary traits and forward and back-transitions (*k* = 6).
2. A full HiSSE model which included a four-level hidden state associated with the binary focal trait, yielding 16 estimates of these parameters plus a single rate for transitions (*k* = 17).

Following the selection of the best-fit model (see below), we calculated diversification rates associated with each of the focal traits. This model configuration is illustrated in **Figure 2**, which shows how the above models increase in complexity and which CID models correspond to which character-dependent models.

**Figure 2.**
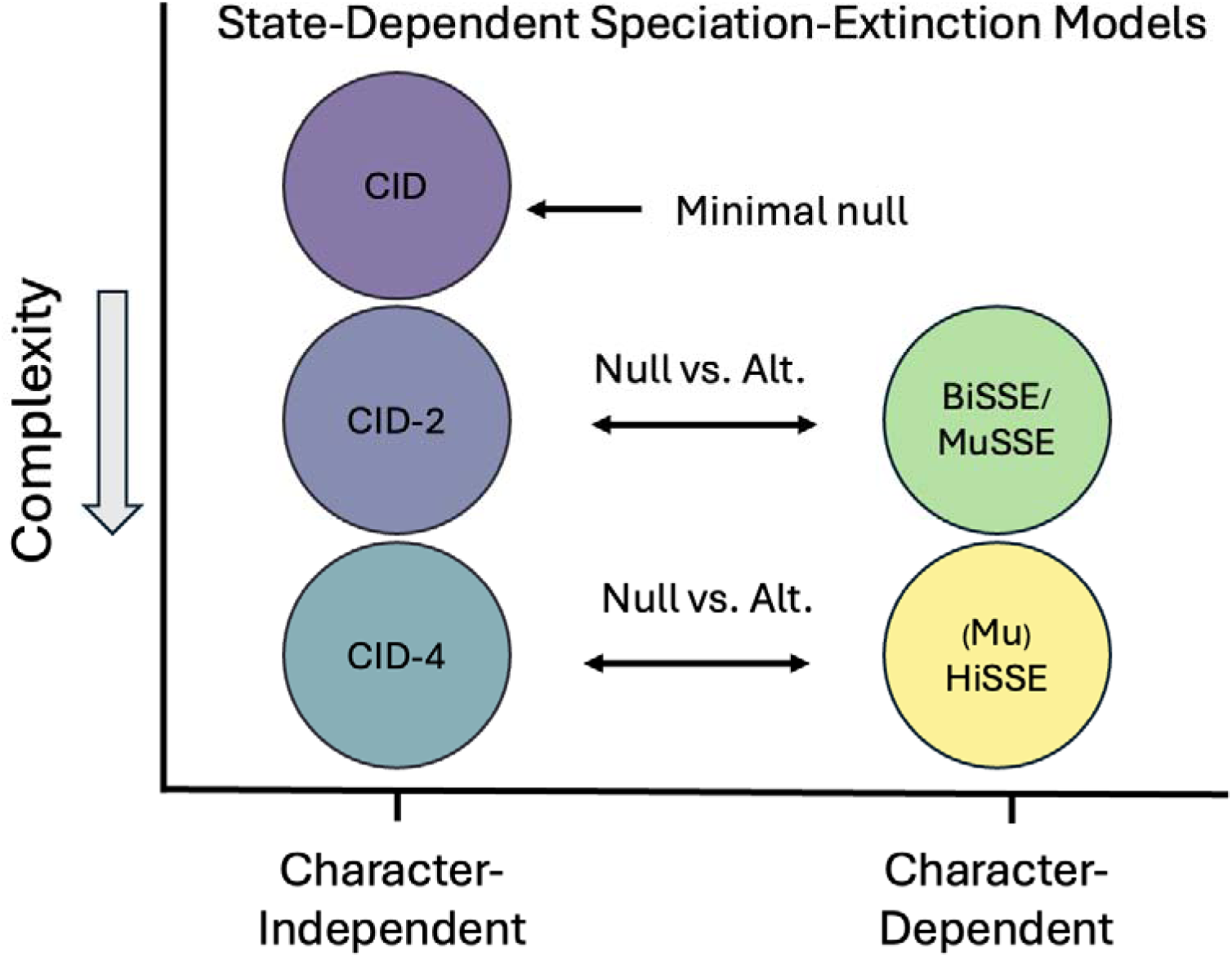
Models for state-dependent speciation-extinction models, in descending order of complexity based on number of parameters. Character-independent diversification models are shown alongside their character-dependent counterparts.

### Analysis of association between sexual system states and diversification

Since we used a three-state classification system for plant sexual systems (dioecy, monoecy, and hermaphroditism; Wang et al. 2021), we utilized the MuHiSSE framework (Multicharacter Hidden State Speciation and Extinction; Nakov et al. 2018), an expansion of HiSSE which allows for traits with more than two states. Utilizing this framework also has the advantage of being directly comparable to other results obtained from HiSSE (Beaulieu and O’Meara, 2016). MuHiSSE was initially designed for two pairs of correlated traits (i.e., 4 states). However, implementation of MuHiSSE with three-character states is also possible and is documented for `hisse` 2.1.13 by the package authors (see: https://speciationextinction.info/). We used the MuHiSSE functionality following the same basic set of models as with HiSSE, with a few modifications to avoid over-parameterization of the more complex multi-state models. These are as follows:

1. A minimal null, CID, with speciation and extinction parameters set to be equal for all three traits and four variable transition rates (*k* = 6).
2. A character-independent model, CID-2, designed to be the null equivalent in complexity to a standard MuSSE model (i.e., multi-character but no hidden states), with equal rates of speciation/extinction for all three states and fixed transition rates between states (*k =* 5).
3. A character-independent model, CID-4, designed to be the null equivalent of a three-state HiSSE model with correlated hidden states; here extinction rate and transitions are fixed, and speciation rate is allowed to vary (*k* = 6).
4. A character-dependent model with no hidden states (MuSSE) which includes the three sexual system traits but does not include any hidden states, with extinction rate fixed, and speciation rate allowed to vary (*k* = 8).
5. A full character-dependent hidden state (HiSSE) model, which includes three-state trait data and a correlated hidden state, with varying speciation and fixed extinction, and in which transitions between states are allowed to vary (*k =* 16).
6. A full character-dependent hidden state model (Full-8) which includes three-state trait data and a correlated hidden state, allowing for both varying diversification and extinction (i.e., eight parameters from speciation and extinction), and variation in transitions between states (*k =* 21).

Since the HiSSE framework calculates extinction fraction and rate of turnover (Beaulieu and O’Meara, 2016), we also re-parameterize these values into speciation and extinction rates using a custom function to back-transform from extinction fraction and turnover (Revell and Harmon, 2022). Then the rate of net diversification was calculated directly from speciation and extinction rates (λ – μ). The 95% confidence intervals associated with each inferred parameter were calculated using adaptive sampling of the likelihood surface.

### Model selection and testing for cross-family patterns

For both binary-state and three-state data, the best-fit model was chosen using AICc (Hurvich and Tsai, 1989), a corrected version of AIC (Akaike 1974) which is less prone to overfitting. We also calculated the Akaike weights (Wagenmakers and Farrell 2004). Model averaging was employed in cases in which ΔAIC was less than two.

We tested the general hypothesis that our focal traits which reduce self-fertilization (i.e., dioecy and monoecy) are associated with higher rates of diversification in two ways. First, we assessed the consistency of patterns across families using a chi-square test, where we tested whether the trait states expected to have the highest net diversification rates (dioecy and monoecy) differed from expectation. We tested against multiple expected values for the distribution: 1) a random distribution across the three sexual system character traits, and 2) a distribution corresponding to overall frequency of the sexual system states across the overall sexual system dataset (Wang et al. 2021). This was not conducted for mating system for two reasons: an overly small sample size (*n* = 3) and a lack of character-dependence.

Second, we used a one-way ANOVA with randomization to look at the magnitude of the relationship between net diversification rates and traits in families where character state impact was supported by our best-fit models. Given the non-independence of estimates within families and the conditionally independent relationships of the families given the mega-phylogeny from Smith and Brown (2018), we used a standard nonparametric randomization approach to generate the null distribution of the *F*-ratio. Data (diversification rate averaged across hidden states) was randomized across sexual systems within families, and values of *F* were calculated iteratively across 9,999 replications to create a null distribution. The value of *F* generated by the non-randomized data was then compared to the frequency distribution generated using the randomized data.

## Results

### Family Trait-Level Coverage

Two out of the 18 families had sufficient data for both mating and sexual system analysis (Asteraceae and Solanaceae). The number of species for which tip data for either mating system or sexual system was available varied considerably between different family groups, with the smallest number of tips corresponding to our minimum threshold of 90 (Rutaceae and Vitaceae) to a maximum of 2,214 (Asteraceae). Percent cover also varied from a low of 1.93% (Asteraceae mating system analysis) to a high of 22.28% (Cucurbitaceae sexual system analysis), with an overall average of 10.16% and a median of 9.10%. The average number of tips on a family-level tree was 419, and the median number of tips was 226. Overall, for the purposes of our cross-family analysis, our priority was to include the maximum number of families possible while following suggested best practices (Maddison et al. 2007, Gamisch 2016).

The families for which we had sufficient tip data for use with SSE models were well-distributed across the mega-phylogeny from Smith and Brown (2018). This is shown in **Figure 3**, which is a version of the Smith and Brown (2018) tree with tips collapsed by a factor of 10, and annotated to show where our tip data falls on the tree. However, we do note that some portions of the tree have noticeable gaps. This was most pronounced in two areas: one corresponds to the orders Asparagales, Myrtales, Saxifragales, and Ranunculales; the second corresponds to the orders Caryophyllales, Ericales, and Gentianales. Despite these gaps in coverage, to the best of our knowledge this is the most comprehensive study which addresses plant mating and sexual systems with SSE models to date.

**Figure 3.**
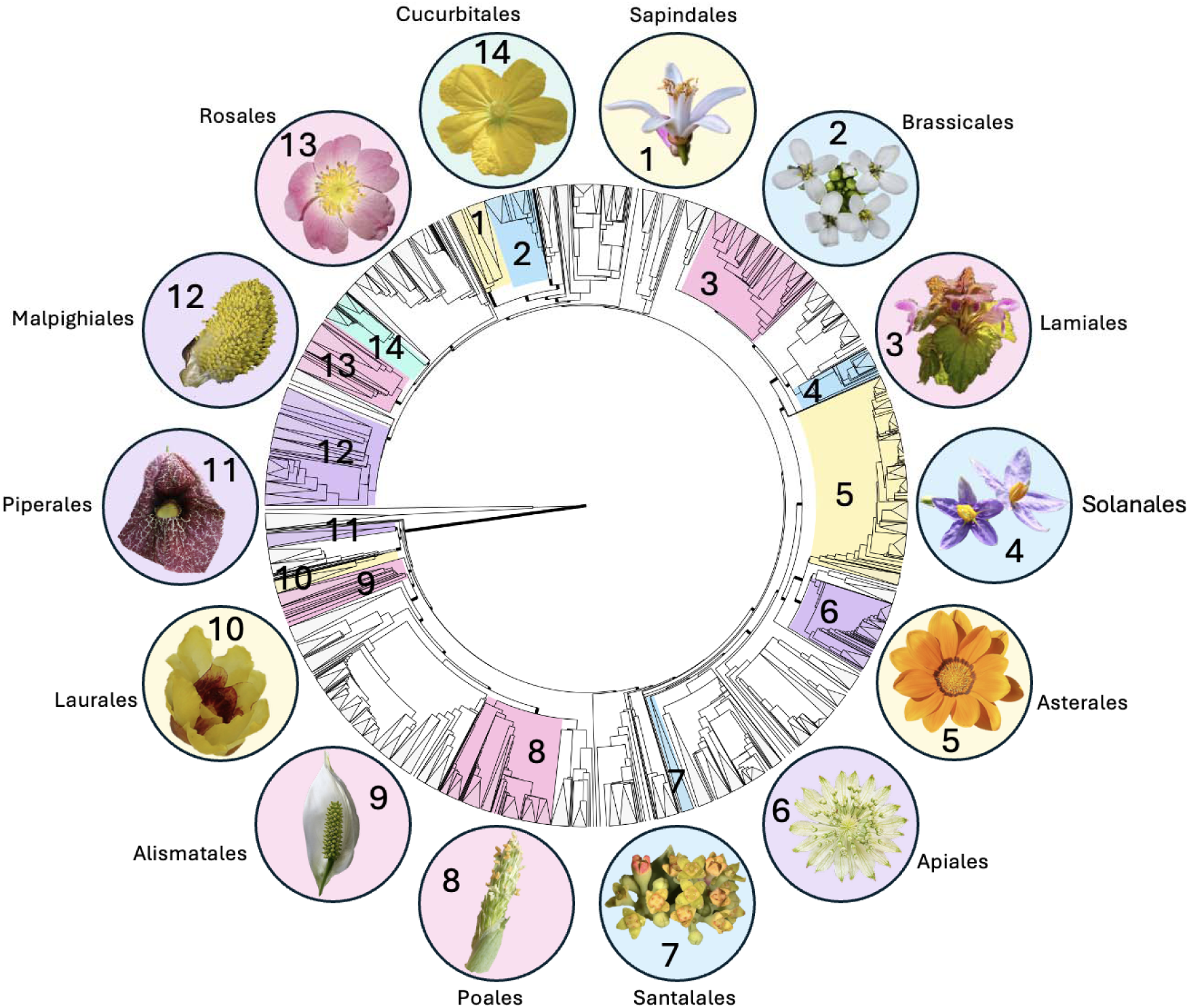
Phylogeny modified from Smith and Brown (2018), showing the distribution of the fourteen angiosperm orders which have at least one representative family in this analysis. Tips are collapsed by a factor of 10 to allow for better visualization. Attributions and identifications for the example species for each order can be found in **Table S1.**

### Families with Mating System Data

We had a limited number of families with sufficient mating system coverage at tips for analysis with SSE methods (*n* = 3, Asteraceae, Brassicaceae, and Solanaceae). Onagraceae, while previously analyzed elsewhere (Freyman and Höhna, 2019), did not have sufficient matches between species with known trait data and the Smith and Brown (2018) tree, and was omitted for consistency with the remainder of the dataset.

Mating system was not a better predictor of rates of diversification than character-independent speciation and extinction across all three of these families using HiSSE (Table 2). The best-fit model for Asteraceae was CID-4 (ΔAICc = 21.88, *ω_i_* = 1.00), for Brassicaceae CID- 2 (ΔAIC = 11.04, *ω_i_* = 0.98), and for Solanaceae CID-4 (ΔAICc = 4.62; *ω_i_* = 0.82). The above models were better supported in all cases than the full hidden-state models, which were the next-best supported model in all cases. However, it is notable that for Solanaceae the difference between the best-fit model (CID-4) and the next-best-fit model (full) is less pronounced than for either Asteraceae or Brassicaceae. Although the number of hidden non-mating system states which best described the families varied (with a simpler model better explaining results in Brassicaceae), these results consistently point to mating system not being a strong predictor of diversification rate.

**Table 2.** Comparison of character-dependent and character-independent models for mating system (i.e., a binary: selfing or non-selfing) using HiSSE. Models which are supported by AIC are shown in bold.

|  | **model** | **logL** | **K** | **AIC** | **AICc** | **Akaike.weight** |
| --- | --- | --- | --- | --- | --- | --- |
| **Asteraceae** | CID | -1302.02 | 4 | 2612.04 | 2612.13 | 0.00 |
|  | BiSSE | -1291.55 | 6 | 2595.09 | 2595.27 | 0.00 |
|  | CID-2 | -1225.73 | 5 | 2461.47 | 2461.59 | 0.00 |
|  | **CID-4** | **-1197.38** | **9** | **2412.76** | **2413.15** | **1.00** |
|  | Full | -1199.65 | 17 | 2433.31 | 2434.64 | 0.00 |
| **Brassicaceae** | CID | -796.08 | 4 | 1600.15 | 1600.36 | 0.00 |
|  | BiSSE | -760.63 | 6 | 1533.26 | 1533.69 | 0.00 |
|  | **CID-2** | **-734.52** | **5** | **1479.04** | **1479.35** | **0.98** |
|  | CID-4 | -735.56 | 9 | 1489.13 | 1490.08 | 0.01 |
|  | Full | -726.55 | 17 | 1487.11 | 1490.47 | 0.02 |
| **Solanaceae** | CID | -868.84 | 4 | 1745.68 | 1745.83 | 0.00 |
|  | BiSSE | -861.29 | 6 | 1734.59 | 1734.89 | 0.00 |
|  | CID-2 | -841.14 | 5 | 1692.27 | 1692.49 | 0.00 |
|  | **CID-4** | **-830.81** | **9** | **1679.63** | **1680.28** | **0.82** |
|  | Full | -824.30 | 17 | 1682.60 | 1684.90 | 0.18 |

The full set of values used for model selection in families analyzed with mating system as the trait of interest are given in **Table 2**. We also present values for turnover and extinction fraction as natively calculated by HiSSE (Beaulieu and O’Meara 2016), as well as the calculated values for speciation, extinction, and net diversification rates in **Table 3**. The values for net diversification ranged between a minimum of -4.16 and a maximum of 0.44. For these values, because character-independent models were determined to be the best-fit, values associated with selfing and outcrossing are held equal. Additionally, according to best practices (Boyko and O’Meara 2024), we provide the 95% confidence intervals associated with these estimates in Table S2.

**Table 3.** Inferred diversification parameters from mating system analysis with a binary-state trait. Columns refer to speciation (λ), extinction (μ), turnover (λ + μ), extinction fraction (μ / λ), and net diversification (net. div.) (λ - μ). Note that for CID models, state 0 is set equal to state 1. Thus, results for both states 0 and 1 are displayed on one line of the table. Letters (A-D) refer to the hidden states in the model.

| | Model | State | $\lambda$ | $\mu$ | $\mu + \lambda$ | $\mu / \lambda$ | Net div. |
| --- | --- | --- | --- | --- | --- | --- | --- |
| **Asteraceae** | CID-4 | 0A/1A | 4902.66 | 4906.83 | 9809.49 | 1.00 | -4.16 |
|  | CID-4 | 0B/1B | 0.00 | 0.00 | 0.00 | 2.58 | 0.00 |
|  | CID-4 | 0C/1C | 9.80 | 9.46 | 19.26 | 0.97 | 0.34 |
|  | CID-4 | 0D/1D | 109.36 | 108.92 | 218.28 | 1.00 | 0.44 |
| **Brassicaceae** | CID-2 | 0A/1A | 0.24 | 0.00 | 0.24 | 0.00 | 0.24 |
|  | CID-2 | 0B/1B | 12.50 | 12.51 | 25.02 | 1.00 | -0.01 |
| **Solanaceae** | CID-4 | 0A/1A | 4.40 | 4.03 | 8.44 | 0.92 | 0.37 |
|  | CID-4 | 0B/1B | 0.09 | 0.00 | 0.09 | 0.00 | 0.09 |
|  | CID-4 | 0C/1C | 1.03 | 0.70 | 1.73 | 0.68 | 0.33 |
|  | CID-4 | 0D/1D | 0.09 | 0.00 | 0.09 | 0.00 | 0.09 |

### Families with Sexual System Data

We obtained 17 families with sufficient data for analysis using sexual system data. This count includes the two families mentioned above which had both mating system and sexual system data (Asteraceae and Solanaceae). Out of these families, four out of the seventeen only had two states for sexual system present on the trimmed tree (Cucurbitaceae, Lamiaceae, Moraceae, and Vitaceae). This was confirmed to be a product of the true underlying reproductive biology of these families, rather than an artifact of the tree trimming process: all these families exhibited only two sexual systems based on the review in Wang et al. (2021). Specifically, for Cucurbitaceae, Lamiaceae, and Moraceae there was no hermaphroditism (with a single-species exception: *Ajuga reptans* in the Lamiaceae family; however, this species did not appear on our trait-matched tree) and in Vitaceae there was no monoecy. For these families with two states, model selection values are given in **Table 4**, and values deriving from the best-fit SSE model are given in **Table 5**. The remaining thirteen families had tips with all three states, with model selection detailed in **Table 6** and values for each of the three sexual system states in **Table 7**.

**Table 4.** Comparison for character-dependent and character-independent models for sexual systems (i.e., dioecious, monoecious, or hermaphrodite) which have two states using HiSSE. Models which are best supported as determined by AICc are shown in bold. An asterisk indicates families for which model averaging was used in the analysis.

|  | **Model** | **logL** | **k** | **AIC** | **AICc** | **Akaike.weight** |
| --- | --- | --- | --- | --- | --- | --- |
| **Cucurbitaceae** | CID | -666.40 | 4 | 1340.81 | 1341.00 | 0.00 |
|  | BiSSE | -666.27 | 6 | 1344.54 | 1344.95 | 0.00 |
|  | CID-2 | -654.95 | 5 | 1319.90 | 1320.19 | 0.00 |
|  | CID-4 | -640.40 | 9 | 1298.80 | 1299.68 | 0.04 |
|  | **Full** | **-629.29** | **17** | **1292.59** | **1295.70** | **0.96** |
| **Lamiaceae** | CID | -1326.04 | 4 | 2660.08 | 2660.16 | 0.00 |
|  | BiSSE | -1196.06 | 6 | 2404.13 | 2404.30 | 0.00 |
|  | CID-2 | -1308.50 | 5 | 2626.99 | 2627.12 | 0.00 |
|  | CID-4 | -1298.98 | 9 | 2615.96 | 2616.34 | 0.00 |
|  | **Full** | **-1145.06** | **17** | **2324.12** | **2325.42** | **1.00** |
| **Moraceae\*** | CID | -617.53 | 4 | 1243.06 | 1243.23 | 0.00 |
|  | BiSSE | -615.76 | 6 | 1243.52 | 1243.90 | 0.00 |
|  | CID-2 | -574.27 | 5 | 1158.53 | 1158.80 | 0.01 |
|  | **CID-4** | **-566.65** | **9** | **1151.30** | **1152.12** | **0.41** |
|  | **Full** | -558.29 | 17 | 1150.57 | 1153.43 | 0.58 |
| **Vitaceae** | CID | -368.81 | 4 | 745.62 | 746.09 | 0.00 |
|  | BiSSE | -367.15 | 6 | 746.30 | 747.31 | 0.00 |
|  | **CID-2** | **-343.54** | **5** | **697.08** | **697.80** | **0.97** |
|  | CID-4 | -343.19 | 9 | 704.37 | 706.62 | 0.03 |
|  | Full | -336.98 | 17 | 707.96 | 716.46 | 0.00 |

**Table 5.** Inferred diversification parameters from sexual system analysis in families with two observed sexual system states. Columns refer to speciation (λ), extinction (μ), turnover (λ + μ), extinction fraction (μ / λ), and net diversification (net. div.) (λ - μ). Letters (A-D) refer to the hidden states in the model.

| | Model | State | $\lambda$ | $\mu$ | $\lambda + \mu$ | $\mu / \lambda$ | Net div. |
| --- | --- | --- | --- | --- | --- | --- | --- |
| **Cucurbitaceae** | Full | **0A** | 0.10 | 0.00 | 0.10 | 0.00 | 0.10 |
|  | Full | **1A** | 23.80 | 23.70 | 47.50 | 1.00 | 0.06 |
|  | Full | **0B** | 0.53 | 0.35 | 0.88 | 0.65 | 0.19 |
|  | Full | **1B** | 2570.00 | 7430.00 | 10000.00 | 2.89 | -4860.00 |
|  | Full | **0C** | 4.09 | 3.78 | 7.88 | 0.92 | 0.31 |
|  | Full | **1C** | 0.73 | 0.37 | 1.10 | 0.51 | 0.36 |
|  | Full | **0D** | 0.09 | 0.00 | 0.09 | 0.00 | 0.09 |
|  | Full | **1D** | 0.10 | 0.18 | 0.28 | 1.87 | -0.08 |
| **Lamiaceae** | Full | **0A** | 2480.00 | 7410.00 | 9890.00 | 2.99 | -4930.00 |
|  | Full | **1A** | 3.33 | 0.00 | 3.33 | 0.00 | 3.33 |
|  | Full | **0B** | 50.80 | 152.00 | 203.00 | 3.00 | -102.00 |
|  | Full | **1B** | 3.07 | 0.00 | 3.07 | 0.00 | 3.07 |
|  | Full | **0C** | 2490.00 | 7460.00 | 9950.00 | 3.00 | -4970.00 |
|  | Full | **1C** | 3.37 | 0.22 | 3.59 | 0.07 | 3.15 |
|  | Full | **0D** | 0.00 | 0.00 | 0.00 | 2.77 | 0.00 |
|  | Full | **1D** | 4980.00 | 5020.00 | 10000.00 | 1.01 | -30.10 |
| **Moraceae** | CID-4 | **0A** | 0.12 | 0.00 | 0.12 | 0.00 | 0.12 |
|  | CID-4 | **1A** | 0.12 | 0.00 | 0.12 | 0.00 | 0.12 |
|  | CID-4 | **0B** | 0.00 | 0.00 | 0.00 | 1.50 | 0.00 |
|  | CID-4 | **1B** | 0.00 | 0.00 | 0.00 | 1.50 | 0.00 |
|  | CID-4 | **0C** | 0.12 | 0.00 | 0.12 | 0.00 | 0.12 |
|  | CID-4 | **1C** | 0.12 | 0.00 | 0.12 | 0.00 | 0.12 |
|  | CID-4 | **0D** | 2.67 | 2.25 | 4.92 | 0.85 | 0.41 |
|  | CID-4 | **1D** | 2.67 | 2.25 | 4.92 | 0.85 | 0.41 |
| **Vitaceae** | CID-2 | **0A** | 0.11 | 0.00 | 0.11 | 0.00 | 0.11 |
|  | CID-2 | **1A** | 0.11 | 0.00 | 0.11 | 0.00 | 0.11 |
|  | CID-2 | **0B** | 2.37 | 2.12 | 4.49 | 0.90 | 0.24 |
|  | CID-2 | **1B** | 2.37 | 2.12 | 4.49 | 0.90 | 0.24 |

**Table 6.** Comparison for character-dependent and character-independent models for sexual systems (i.e., dioecious, monoecious, or hermaphrodite) which have three states using MuHiSSE. Models which are supported by AIC are shown in bold.

|  | Model | logL | k | AIC | AICc | Akaike.weight |
| --- | --- | --- | --- | --- | --- | --- |
| **Apiaceae** | CID | -1188.47 | 6 | 2388.94 | 2389.17 | 0.00 |
|  | MuSSE | -1125.22 | 8 | 2266.44 | 2266.83 | 0.00 |
|  | CID-2 | -1152.78 | 5 | 2315.56 | 2315.72 | 0.00 |
|  | CID-4 | -1178.54 | 6 | 2369.08 | 2369.31 | 0.00 |
|  | Full | -1060.06 | 16 | 2152.12 | 2153.64 | 0.00 |
|  | **Full.8** | **-1047.10** | **21** | **2136.21** | **2138.82** | **1.00** |
| **Asteraceae** | CID | -4308.77 | 6 | 8629.54 | 8629.58 | 0.00 |
|  | MuSSE | -4274.33 | 8 | 8564.65 | 8564.72 | 0.00 |
|  | CID-2 | -4120.17 | 4 | 8248.34 | 8248.36 | 0.00 |
|  | CID-4 | -3972.57 | 6 | 7957.14 | 7957.18 | 0.00 |
|  | **Full** | **-3607.73** | **16** | **7247.46** | **7247.71** | **0.92** |
|  | Full.8 | -3605.13 | 21 | 7252.25 | 7252.67 | 0.08 |
| **Araceae** | CID | -1109.58 | 6 | 2231.16 | 2231.55 | 0.00 |
|  | MuSSE | -1087.05 | 8 | 2190.10 | 2190.78 | 0.00 |
|  | CID-2 | -1093.05 | 5 | 2196.10 | 2196.38 | 0.00 |
|  | CID-4 | -1095.13 | 6 | 2202.26 | 2202.66 | 0.00 |
|  | Full | -1055.66 | 16 | 2143.33 | 2145.99 | 0.07 |
|  | **Full.8** | **-1048.04** | **21** | **2138.08** | **2142.72** | **0.93** |
| **Euphorbiaceae** | CID | -2266.13 | 6 | 4544.26 | 4544.36 | 0.00 |
|  | MuSSE | -2219.11 | 8 | 4454.22 | 4454.39 | 0.00 |
|  | CID-2 | -2239.39 | 5 | 4488.78 | 4488.85 | 0.00 |
|  | CID-4 | -2138.27 | 6 | 4288.54 | 4288.64 | 0.00 |
|  | Full | -2090.04 | 16 | 4212.09 | 4212.75 | 0.00 |
|  | **Full.8** | **-2061.54** | **21** | **4165.08** | **4166.21** | **1.00** |
| **Lauraceae** | CID | -580.76 | 6 | 1173.53 | 1173.93 | 0.00 |
|  | MuSSE | -563.32 | 8 | 1142.64 | 1143.35 | 0.00 |
|  | CID-2 | -591.41 | 5 | 1192.82 | 1193.11 | 0.00 |
|  | CID-4 | -570.83 | 6 | 1153.65 | 1154.06 | 0.00 |
|  | **Full** | **-527.42** | **16** | **1086.84** | **1089.60** | **1.00** |
|  | Full.8 | -527.71 | 21 | 1097.43 | 1102.24 | 0.01 |
| **Loranthaceae** | CID | -358.60 | 6 | 729.19 | 730.13 | 0.25 |
|  | MuSSE | -356.76 | 8 | 729.51 | 731.15 | 0.21 |
|  | CID-2 | -369.22 | 5 | 748.44 | 749.09 | 0.00 |
|  | **CID-4** | **-357.91** | **6** | **727.82** | **728.76** | **0.49** |
|  | Full | -350.39 | 16 | 732.78 | 739.58 | 0.04 |
|  | Full.8 | -347.99 | 21 | 737.97 | 750.29 | 0.00 |
| **Piperaceae** | CID | -359.76 | 6 | 731.51 | 732.37 | 0.00 |
|  | MuSSE | -355.41 | 8 | 726.83 | 728.33 | 0.00 |
|  | CID-2 | -363.67 | 5 | 737.34 | 737.94 | 0.00 |
|  | **CID-4** | **-340.65** | **6** | **693.29** | **694.15** | **1.00** |
|  | Full | -346.30 | 16 | 724.60 | 730.79 | 0.00 |
|  | Full.8 | -341.73 | 21 | 725.46 | 736.60 | 0.00 |
| **Poaceae** | CID | -2246.69 | 6 | 4505.39 | 4505.48 | 0.00 |
|  | MuSSE | -2064.22 | 8 | 4144.45 | 4144.62 | 0.00 |
|  | CID-2 | -2286.08 | 5 | 4582.16 | 4582.23 | 0.00 |
|  | CID-4 | -2266.36 | 6 | 4544.72 | 4544.82 | 0.00 |
|  | **Full** | **-1925.16** | **16** | **3882.33** | **3882.97** | **1.00** |
|  | Full.8 | -2008.11 | 21 | 4058.21 | 4059.31 | 0.00 |
| **Rosaceae** | CID | -2521.88 | 6 | 5055.76 | 5055.89 | 0.00 |
|  | MuSSE | -2155.44 | 8 | 4326.87 | 4327.10 | 0.00 |
|  | CID-2 | -2278.26 | 5 | 4566.53 | 4566.62 | 0.00 |
|  | CID-4 | -2306.41 | 6 | 4624.81 | 4624.95 | 0.00 |
|  | **Full** | **-2049.88** | **16** | **4131.76** | **4132.64** | **1.00** |
|  | Full.8 | -2104.90 | 21 | 4251.80 | 4253.29 | 0.00 |
| **Rutaceae** | CID | -360.87 | 6 | 733.75 | 734.76 | 0.00 |
|  | MuSSE | -348.00 | 8 | 712.00 | 713.78 | 0.00 |
|  | CID-2 | -363.37 | 5 | 736.75 | 737.46 | 0.00 |
|  | **CID-4** | **-337.12** | **6** | **686.24** | **687.26** | **0.39** |
|  | Full | -326.75 | 16 | 685.49 | 692.94 | 0.57 |
|  | Full.8 | -324.58 | 21 | 691.15 | 704.74 | 0.03 |
| **Saliaceae** | CID | -292.59 | 6 | 597.17 | 597.74 | 0.00 |
|  | MuSSE | -275.99 | 8 | 567.99 | 568.98 | 0.00 |
|  | CID-2 | -278.35 | 5 | 566.71 | 567.11 | 0.00 |
|  | CID-4 | -189.75 | 6 | 391.51 | 392.08 | 0.00 |
|  | **Full** | **-159.25** | **16** | **350.50** | **354.47** | **1.00** |
|  | Full.8 | -275.20 | 21 | 592.39 | 599.39 | 0.00 |
| **Solanaceae** | CID | -1253.35 | 6 | 2518.70 | 2518.89 | 0.00 |
|  | MuSSE | -1154.12 | 8 | 2324.23 | 2324.54 | 0.00 |
|  | CID-2 | -1223.11 | 5 | 2456.22 | 2456.35 | 0.00 |
|  | CID-4 | -1217.47 | 6 | 2446.94 | 2447.12 | 0.00 |
|  | **Full** | **-1123.53** | **16** | **2279.05** | **2280.25** | **1.00** |
|  | Full.8 | -1128.19 | 21 | 2298.39 | 2300.45 | 0.00 |
| **Urticaceae** | CID | -465.67 | 6 | 943.33 | 944.14 | 0.00 |
|  | MuSSE | -452.97 | 8 | 921.95 | 923.36 | 0.15 |
|  | CID-2 | -557.83 | 5 | 1125.66 | 1126.23 | 0.00 |
|  | CID-4 | -474.04 | 6 | 960.09 | 960.90 | 0.00 |
|  | **Full** | **-443.26** | **16** | **918.52** | **924.30** | **0.85** |
|  | Full.8 | -450.37 | 21 | 942.74 | 953.12 | 0.00 |

**Table 7.**
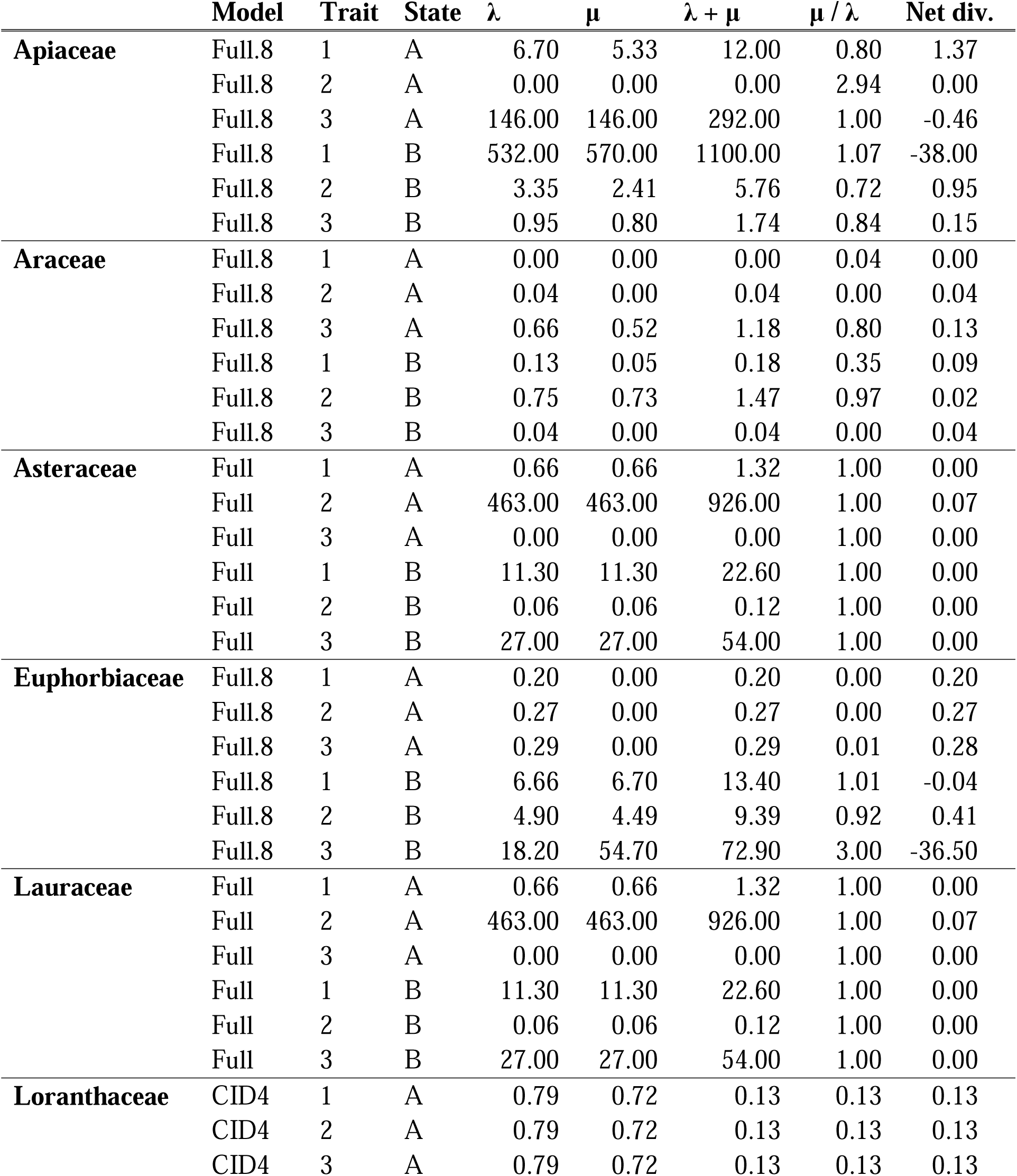
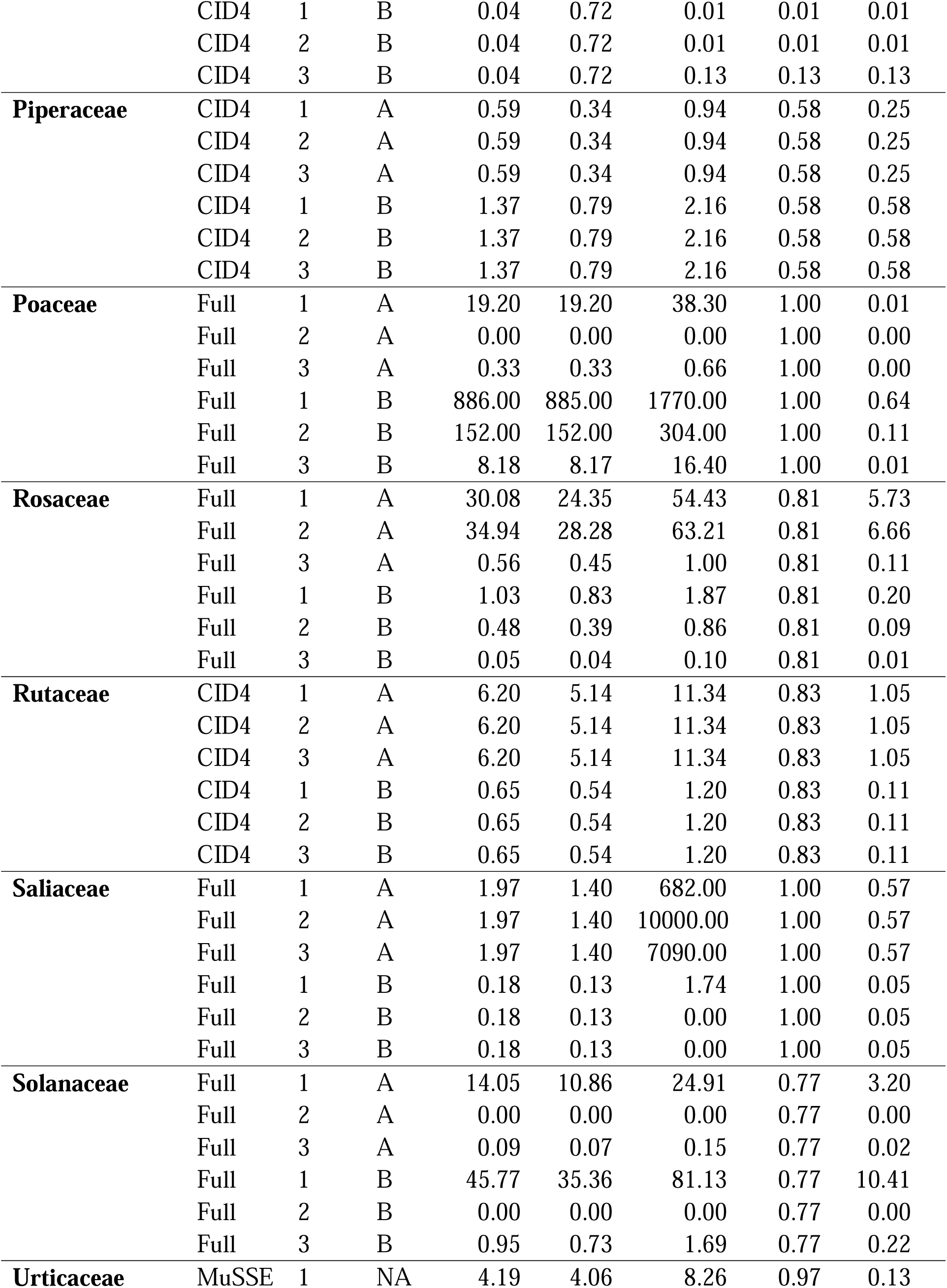
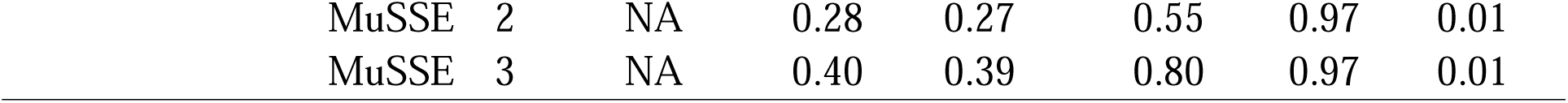
Inferred diversification parameters from sexual system analysis in families with three observed sexual system states. Columns refer to speciation (λ), extinction (μ), turnover (λ + μ), extinction fraction (μ / λ), and net diversification (net. div.) (λ - μ). Under the “trait” column, one corresponds to dioecy, two corresponds to monoecy, and three corresponds to hermaphroditism. Letters (A and B) refer to the hidden states in the model.

| | Model | Trait | State | $\lambda$ | $\mu$ | $\lambda + \mu$ | $\mu / \lambda$ | Net div. |
| --- | --- | --- | --- | --- | --- | --- | --- | --- |
| **Apiaceae** | Full.8 | 1 | A | 6.70 | 5.33 | 12.00 | 0.80 | 1.37 |
|  | Full.8 | 2 | A | 0.00 | 0.00 | 0.00 | 2.94 | 0.00 |
|  | Full.8 | 3 | A | 146.00 | 146.00 | 292.00 | 1.00 | -0.46 |
|  | Full.8 | 1 | B | 532.00 | 570.00 | 1100.00 | 1.07 | -38.00 |
|  | Full.8 | 2 | B | 3.35 | 2.41 | 5.76 | 0.72 | 0.95 |
|  | Full.8 | 3 | B | 0.95 | 0.80 | 1.74 | 0.84 | 0.15 |
| **Araceae** | Full.8 | 1 | A | 0.00 | 0.00 | 0.00 | 0.04 | 0.00 |
|  | Full.8 | 2 | A | 0.04 | 0.00 | 0.04 | 0.00 | 0.04 |
|  | Full.8 | 3 | A | 0.66 | 0.52 | 1.18 | 0.80 | 0.13 |
|  | Full.8 | 1 | B | 0.13 | 0.05 | 0.18 | 0.35 | 0.09 |
|  | Full.8 | 2 | B | 0.75 | 0.73 | 1.47 | 0.97 | 0.02 |
|  | Full.8 | 3 | B | 0.04 | 0.00 | 0.04 | 0.00 | 0.04 |
| **Asteraceae** | Full | 1 | A | 0.66 | 0.66 | 1.32 | 1.00 | 0.00 |
|  | Full | 2 | A | 463.00 | 463.00 | 926.00 | 1.00 | 0.07 |
|  | Full | 3 | A | 0.00 | 0.00 | 0.00 | 1.00 | 0.00 |
|  | Full | 1 | B | 11.30 | 11.30 | 22.60 | 1.00 | 0.00 |
|  | Full | 2 | B | 0.06 | 0.06 | 0.12 | 1.00 | 0.00 |
|  | Full | 3 | B | 27.00 | 27.00 | 54.00 | 1.00 | 0.00 |
| **Euphorbiaceae** | Full.8 | 1 | A | 0.20 | 0.00 | 0.20 | 0.00 | 0.20 |
|  | Full.8 | 2 | A | 0.27 | 0.00 | 0.27 | 0.00 | 0.27 |
|  | Full.8 | 3 | A | 0.29 | 0.00 | 0.29 | 0.01 | 0.28 |
|  | Full.8 | 1 | B | 6.66 | 6.70 | 13.40 | 1.01 | -0.04 |
|  | Full.8 | 2 | B | 4.90 | 4.49 | 9.39 | 0.92 | 0.41 |
|  | Full.8 | 3 | B | 18.20 | 54.70 | 72.90 | 3.00 | -36.50 |
| **Lauraceae** | Full | 1 | A | 0.66 | 0.66 | 1.32 | 1.00 | 0.00 |
|  | Full | 2 | A | 463.00 | 463.00 | 926.00 | 1.00 | 0.07 |
|  | Full | 3 | A | 0.00 | 0.00 | 0.00 | 1.00 | 0.00 |
|  | Full | 1 | B | 11.30 | 11.30 | 22.60 | 1.00 | 0.00 |
|  | Full | 2 | B | 0.06 | 0.06 | 0.12 | 1.00 | 0.00 |
|  | Full | 3 | B | 27.00 | 27.00 | 54.00 | 1.00 | 0.00 |
| **Loranthaceae** | CID4 | 1 | A | 0.79 | 0.72 | 0.13 | 0.13 | 0.13 |
|  | CID4 | 2 | A | 0.79 | 0.72 | 0.13 | 0.13 | 0.13 |
|  | CID4 | 3 | A | 0.79 | 0.72 | 0.13 | 0.13 | 0.13 |
|  | CID4 | 1 | B | 0.04 | 0.72 | 0.01 | 0.01 | 0.01 |
|  | CID4 | 2 | B | 0.04 | 0.72 | 0.01 | 0.01 | 0.01 |
|  | CID4 | 3 | B | 0.04 | 0.72 | 0.13 | 0.13 | 0.13 |
| **Piperaceae** | CID4 | 1 | A | 0.59 | 0.34 | 0.94 | 0.58 | 0.25 |
|  | CID4 | 2 | A | 0.59 | 0.34 | 0.94 | 0.58 | 0.25 |
|  | CID4 | 3 | A | 0.59 | 0.34 | 0.94 | 0.58 | 0.25 |
|  | CID4 | 1 | B | 1.37 | 0.79 | 2.16 | 0.58 | 0.58 |
|  | CID4 | 2 | B | 1.37 | 0.79 | 2.16 | 0.58 | 0.58 |
|  | CID4 | 3 | B | 1.37 | 0.79 | 2.16 | 0.58 | 0.58 |
| **Poaceae** | Full | 1 | A | 19.20 | 19.20 | 38.30 | 1.00 | 0.01 |
|  | Full | 2 | A | 0.00 | 0.00 | 0.00 | 1.00 | 0.00 |
|  | Full | 3 | A | 0.33 | 0.33 | 0.66 | 1.00 | 0.00 |
|  | Full | 1 | B | 886.00 | 885.00 | 1770.00 | 1.00 | 0.64 |
|  | Full | 2 | B | 152.00 | 152.00 | 304.00 | 1.00 | 0.11 |
|  | Full | 3 | B | 8.18 | 8.17 | 16.40 | 1.00 | 0.01 |
| **Rosaceae** | Full | 1 | A | 30.08 | 24.35 | 54.43 | 0.81 | 5.73 |
|  | Full | 2 | A | 34.94 | 28.28 | 63.21 | 0.81 | 6.66 |
|  | Full | 3 | A | 0.56 | 0.45 | 1.00 | 0.81 | 0.11 |
|  | Full | 1 | B | 1.03 | 0.83 | 1.87 | 0.81 | 0.20 |
|  | Full | 2 | B | 0.48 | 0.39 | 0.86 | 0.81 | 0.09 |
|  | Full | 3 | B | 0.05 | 0.04 | 0.10 | 0.81 | 0.01 |
| **Rutaceae** | CID4 | 1 | A | 6.20 | 5.14 | 11.34 | 0.83 | 1.05 |
|  | CID4 | 2 | A | 6.20 | 5.14 | 11.34 | 0.83 | 1.05 |
|  | CID4 | 3 | A | 6.20 | 5.14 | 11.34 | 0.83 | 1.05 |
|  | CID4 | 1 | B | 0.65 | 0.54 | 1.20 | 0.83 | 0.11 |
|  | CID4 | 2 | B | 0.65 | 0.54 | 1.20 | 0.83 | 0.11 |
|  | CID4 | 3 | B | 0.65 | 0.54 | 1.20 | 0.83 | 0.11 |
| **Saliaceae** | Full | 1 | A | 1.97 | 1.40 | 682.00 | 1.00 | 0.57 |
|  | Full | 2 | A | 1.97 | 1.40 | 10000.00 | 1.00 | 0.57 |
|  | Full | 3 | A | 1.97 | 1.40 | 7090.00 | 1.00 | 0.57 |
|  | Full | 1 | B | 0.18 | 0.13 | 1.74 | 1.00 | 0.05 |
|  | Full | 2 | B | 0.18 | 0.13 | 0.00 | 1.00 | 0.05 |
|  | Full | 3 | B | 0.18 | 0.13 | 0.00 | 1.00 | 0.05 |
| **Solanaceae** | Full | 1 | A | 14.05 | 10.86 | 24.91 | 0.77 | 3.20 |
|  | Full | 2 | A | 0.00 | 0.00 | 0.00 | 0.77 | 0.00 |
|  | Full | 3 | A | 0.09 | 0.07 | 0.15 | 0.77 | 0.02 |
|  | Full | 1 | B | 45.77 | 35.36 | 81.13 | 0.77 | 10.41 |
|  | Full | 2 | B | 0.00 | 0.00 | 0.00 | 0.77 | 0.00 |
|  | Full | 3 | B | 0.95 | 0.73 | 1.69 | 0.77 | 0.22 |
| **Urticaceae** | MuSSE | 1 | NA | 4.19 | 4.06 | 8.26 | 0.97 | 0.13 |
|  | MuSSE | 2 | NA | 0.28 | 0.27 | 0.55 | 0.97 | 0.01 |
|  | MuSSE | 3 | NA | 0.40 | 0.39 | 0.80 | 0.97 | 0.01 |

From these results, we found that sexual system explains diversification patterns better than the null models in 13 of the 17 families. In these 13 families, the ‘full’ HiSSE model was best supported, meaning that the models which incorporated sexual system better explained patterns of diversification than character-independent models (*ω_i_* ranged from 0.31 to 1.0 across all models, with only three families with a *ω_i_* < 0.8.). The families which did not follow this pattern, and for which character-independent models better explained patterns of diversification were Loranthaceae (CID-4, *ω_i_*= 0.49), Piperaceae (CID-4, *ω_i_*= 1.00), Rutaceae (CID-4, *ω_i_*= 0.39) and Vitaceae (CID-2, *ω_i_*= 0.96). It is worth noting that these four families are also the smallest tip-matched trees we included in our analysis (*n* = 90-105), and it is possible that we have insufficient power to appropriately detect the effect of the measured trait using these SSE models for these families. Furthermore, support for the null models was more mixed for these families than those for which the “full” model was the best fit, especially in Loranthaceae, Moraceae, and Rutaceae. This could suggest some level of rate heterogeneity, or insufficient model power based on the number of trait-matched tips. However, despite these hints of potential rate heterogeneity, the only family with a null model within the set ΔAICc threshold (*Δ*AIC <2 between models) was Moraceae, for which we conducted model averaging between the two best-fit models (CID-4 and the full model).

Additionally, while the vast majority of these models generated estimates for net diversification which fell in the range of -1 to 1, two families had estimates that were clearly outliers: Lamiaceae (with net diversification rates as low as -4970 in the case of state 0A) and Cucurbitaceae (with net diversification values as low as -4860 in the case of state 1B). The values associated with these families are shown in the tables but were excluded from downstream analyses summarizing overall trends across families. We believe these values are rather artifacts of the tree topology relative to the distribution of trait data, and do not correspond to reliable estimates. For Lamiaceae, the majority of trait variation appears inadequately dispersed within the tree, with almost all rate variation occurring in one small section of the overall topology with small branch lengths. For Cucurbitaceae, values appear reasonably distributed, and the outlying estimates only correspond to state 1B (**Table 5**). Phylogenies showing the trait distribution for both families can be found in the repository for this project (https://github.com/elenacicada/diversification_reproduction).

When examined across families, the sexual system state associated with the highest rate of diversification was heterogeneous. This can be seen in **Figure 4**, which shows the average net diversification as associated with each of the three traits for sexual systems. In total, hermaphroditism has the highest rate of net diversification in only one out of the 11 cases (Figure 4), whereas monoecy had the highest rate in seven out of 11 cases, and dioecy had the highest rate in three out of 11 cases. The estimates for the points displayed in Figure 4 are also given numerically in **Table 5** and **Table 7**, with 95% confidence intervals for all sexual system analyses provided in **Tables S3** and **S4.** The magnitude of the values was mostly constrained, with an overall mean of -0.7796 across all three states.

**Figure 4.**
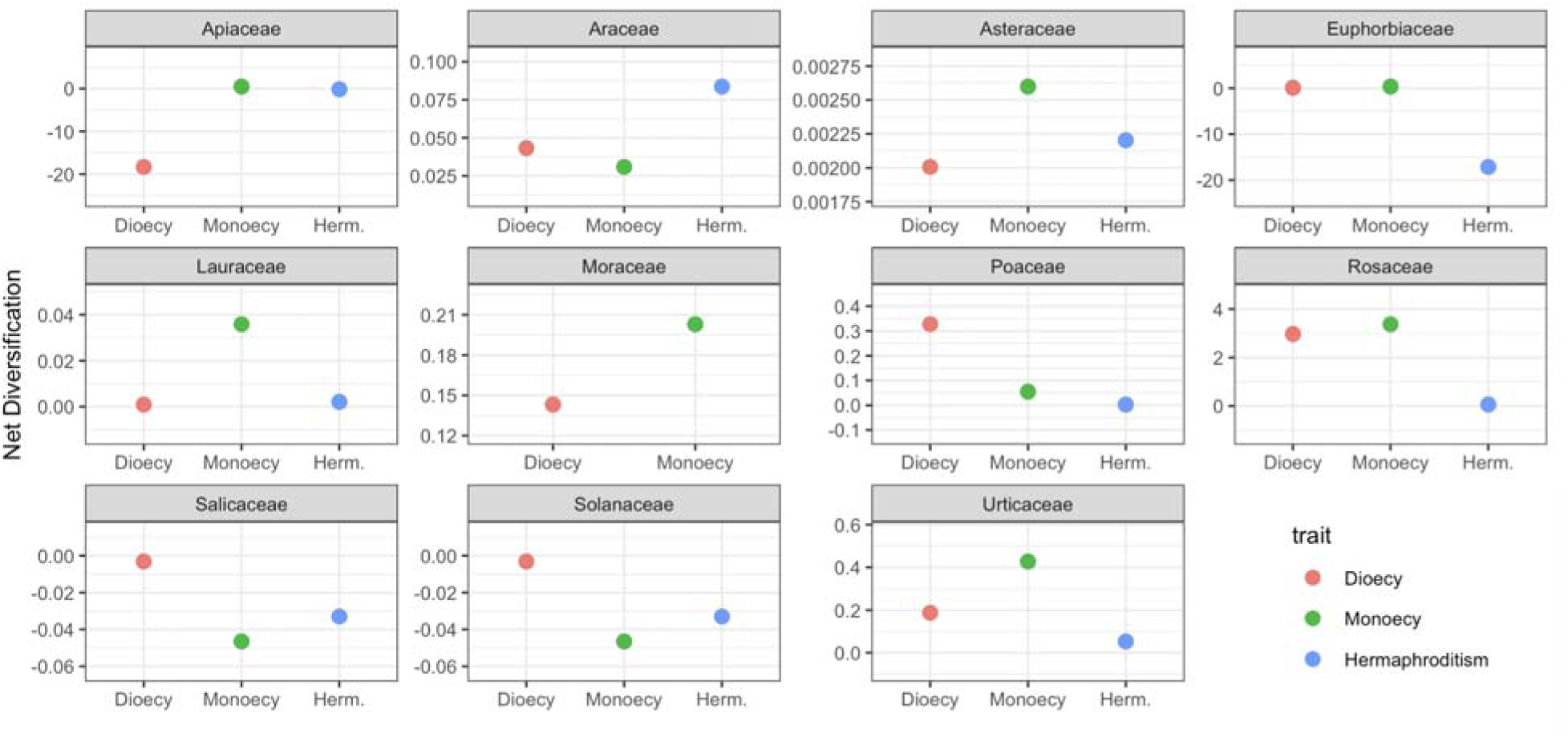
A faceted dot plot showing the mean net diversification associated with sexual system traits for each family in this study. Here, the y-axis has been allowed to vary between faceted plots to capture any general pattern emerging across differing scales.

We also examined how net diversification rate is associated with sexual system across all families individually, which is illustrated in **Figure 5**. In this figure, we are allowing for the y-axis to vary between family groups to better illustrate between-family scale differences. This shows that dioecy or monoecy is the highest-ranked trait for 10 out of 11 families (although note that hermaphroditism is absent for Moraceae). Means for the three sexual system states are also summarized across families in **Figure 6**.

**Figure 5.**
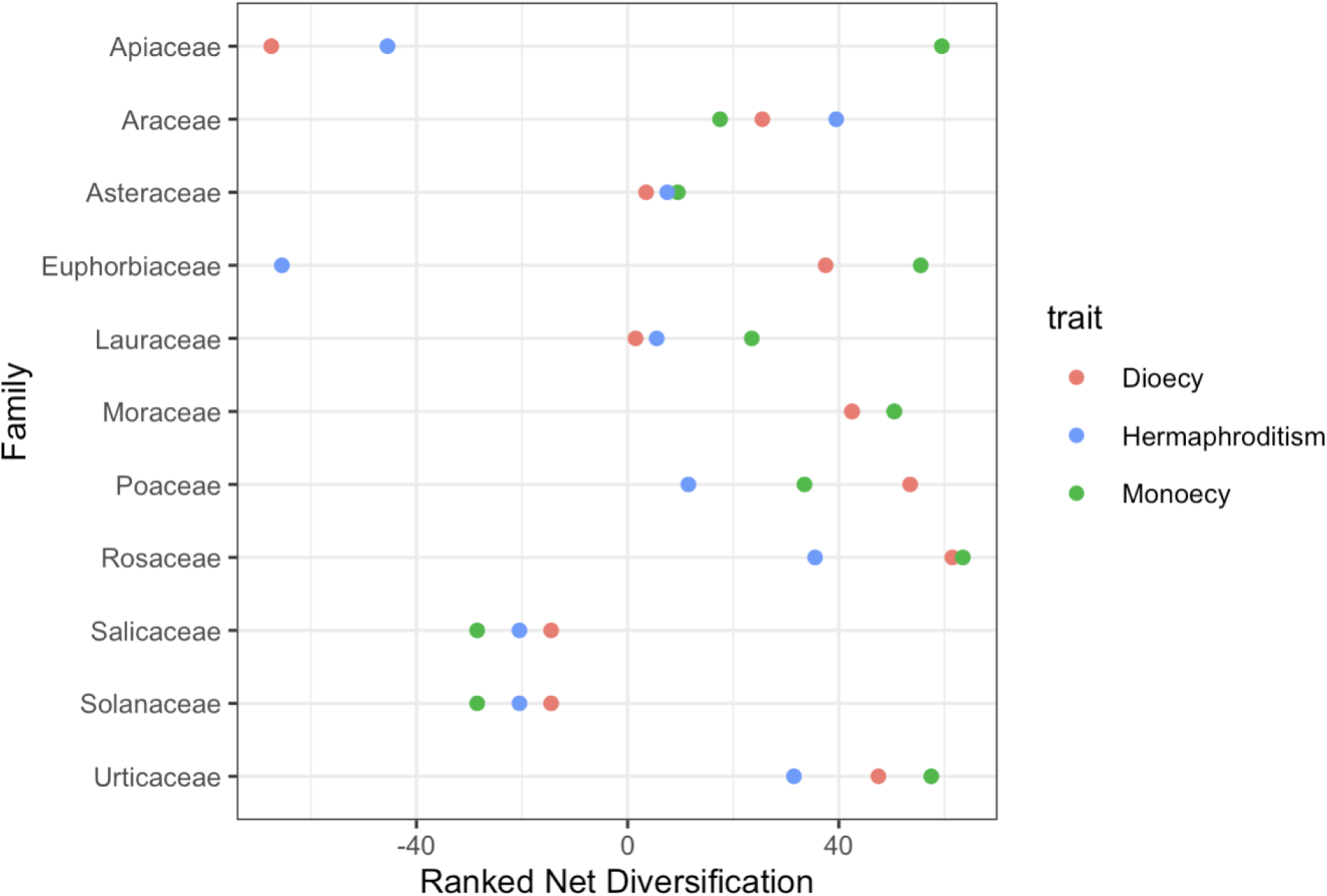
Dot plot showing rates of diversification as associated with sexual system across families. The x-axis shows the (sign-aware) ranked mean of net diversification associated with each trait (averaged across hidden states).

**Figure 6.**
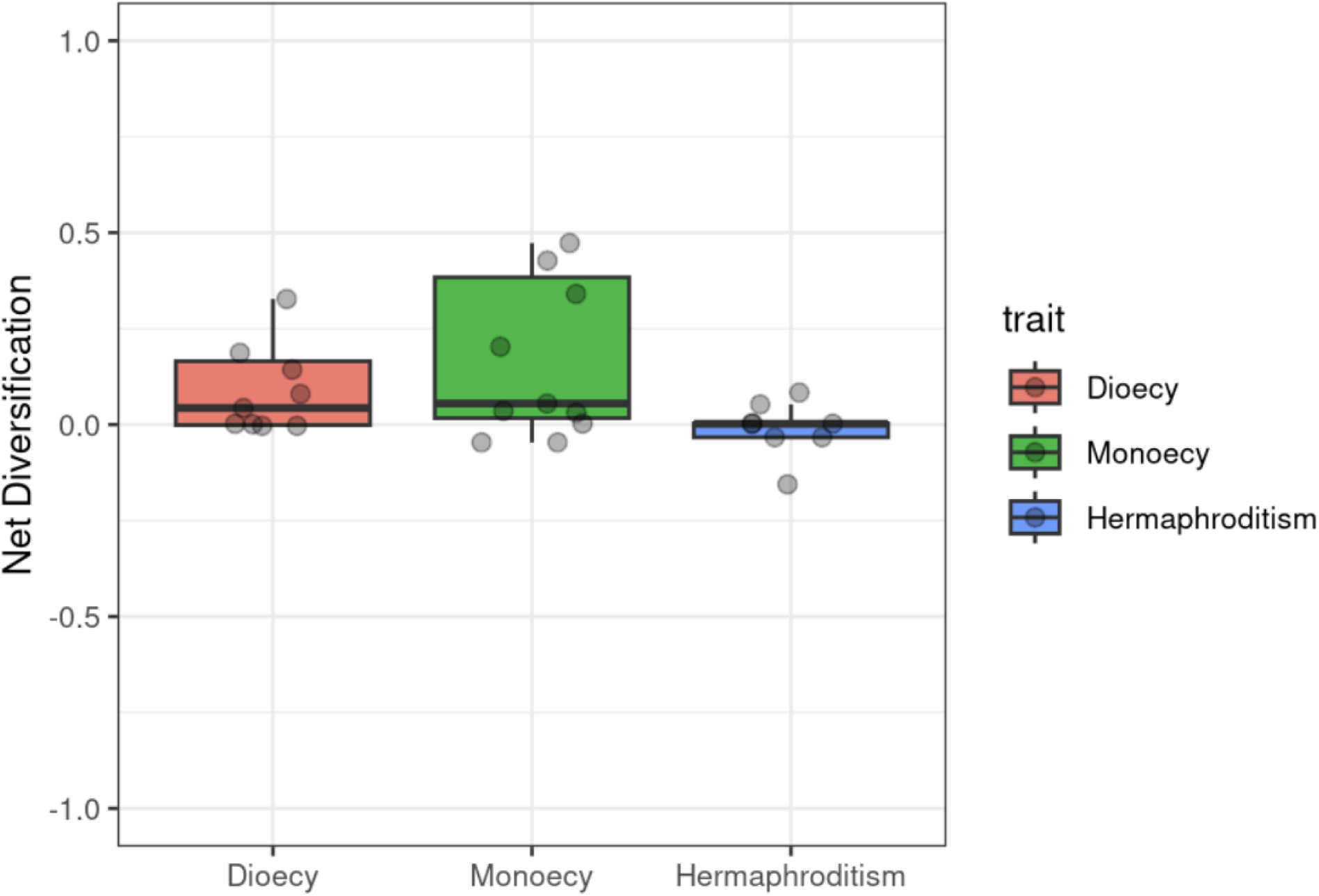
A box-and-whisker plot showing the overall mean net diversification for all families for each sexual system trait. Note for this figure that outlier points shown in Figures 4 and 5 have not been displayed, and the y-axis was limited to +/- 1 to more clearly show means.

We tested for consistency of pattern using a series of chi-squared tests, for which we assumed a null hypothesis of equal counts across families. We looked at 1) the pooled number of highest ranks of diversification rate for monoecy and dioecy compared with hermaphroditism, and 2) all three states separately. These results were not significant, but may indicate a trend. The result for the pooled chi-squared test had a p-value of 0.088 (χ_2_= 2.909, df = 1); and the result for the three-level test had a p-value of 0.078 ( = 5.091, df = 2). Because our sample size was small, we also conducted a binomial test on the pooled data, obtaining a p-value of 1.694e-05.

We also conducted an ANOVA with randomization to test for the magnitude of this pattern. The F-value generated by our data was 0.6015, with a non-significant p-value of 0.5547. The results of the randomization yielded a similar p-value of 0.51025.

## Discussion

### Data and Model Scope

Trait-dependent speciation-extinction models have been, since their initial publication (Madison et al. 2007), in a state of ongoing evolution. Following their initial adoption, subsequent studies highlighted major limitations to the typical use of these models. By comparing to a relatively minimal null (the CID model), BiSSE and related methods were prone to a high rate of false discovery and found “significant” results even in simulated data (Rabosky and Goldberg 2015). Hidden-state models (HiSSE, Beaulieu and O’Meara 2016) and non-parametric options (FiSSE, Rabosky and Goldberg 2017) have both since been proposed to address this issue. More recently, synthesis of existing SSE studies on angiosperms has found that these methods are profoundly impacted by factors outside of focal traits such as the underlying phylogenies used and the modeling method from the larger SSE ecosystem used by investigators (Helmstetter et al. 2022). Furthermore, a recent review found that within the field of mating system research, results across studies have been inconsistent (Marie-Orleach et al. 2024), rather than yielding a generalizable pattern. Different families from across different studies have yielded differing rates of speciation, extinction, and thus net diversification associated with mating system. Similar to the findings of Helmstetter et al. (2022) in SSEs, other studies in the broader field corroborate the idea that individual analysis choices, even when using the same dataset, could yield differing results (Gould et al. 2025). With this in mind, it is unclear whether these differences are driven by analysis choices or reflect true consequences of the underlying reproductive biology.

This highlights the importance of one of the aims of this study, which is to provide cross-family analysis with a consistent underlying phylogeny and standardized methods. Toward this aim, we achieved fairly broad sampling across one of the largest available angiosperm macro- phylogenies (Smith and Brown 2018). However, it is worth noting that some of the areas on our tree which were not sampled correspond to orders which contain large and notably under- sampled families with respect to mating systems, such as Rubiaceae and Orchidaceae (Meyer et al. 2025).

### Mating System Patterns

Results from our analyses using mating system traits led to acceptance of the character- independent models with hidden states. These are the equivalent of “null” models, meaning that the inclusion of mating system trait data does not help better explain patterns of speciation and extinction on the phylogeny. This was consistent across all three families analyzed (Asteraceae, Brassicaceae, and Solanaceae). While previous studies of the same families (i.e., Goldberg et al. 2010 in Solanaceae) had found support for trait-dependent models, the addition of hidden state models (Beaulieu and O’Meara, 2016) led to results with character-independent diversification as the best model. Other previous studies, such as Gamisch et al. (2015) in Orchidaceae, failed to conclusively support or reject the extinction-driven dead-end hypothesis, likely due to limited sample size. This is also a possible constraint for our work: for example, in Asteraceae, while our trait-matched tree contained 477 species, this represents a very small portion of the overall diversity of one of the largest angiosperm families.

In Goldberg et al. (2010), the genus *Solanum* within Solanaceae was analyzed for 356 species using BiSSE. Through this work, they found support for a higher rate of speciation associated with self-compatibility compared to self-incompatibility, but this advantage in speciation rates was erased by a high extinction rate, yielding a higher rate of net diversification for self-incompatible linages overall. Indeed, in our analysis, BiSSE was also better supported by AICc than the CID null model which was standard at the time. In other words, prior to the publication and adoption of the HiSSE framework (Beaulieu and O’Meara, 2016), our result would have led us to the same conclusions (i.e., without hidden-state models, the BiSSE model is better than the CID model by a ΔAICc of 10.94). This pattern of initial analysis using SSE models over-inferring trait-driven patterns of speciation and extinction based on methodological limitations has also been noted by other authors (Herrera-Alsina et al. 2019), which suggests that the differences between our conclusions and those of earlier SSE work in mating systems are a likely result of improvements to the SSE framework (i.e., HiSSE; Beaulieu and O’Meara 2016). This highlights how the present study differs from those prior and that applying SSE models systematically across families is helpful towards the goal of showing general patterns in evolutionary dynamics.

It is also noteworthy that CID models continue to be the best-fit models when rates of extinction are allowed to vary (as in the full binary-state models shown in **Table 3** and **Table 5**). In a framework where selfing is associated with both increased speciation and increased extinction, but an overall negative net diversification is driven by high rates of extinction (as in Goldberg et al. 2010), we might imagine that fixing extinction rates could limit our ability to accurately depict the impacts of selfing in our character-dependent models. However, this does not seem to be the case, as allowing extinction to vary does not change our results.

The acceptance of the more-complex null models from the HiSSE framework supports the hypothesis that, while drivers of speciation and extinction exist for these families at varying levels of complexity (as suggested by differing numbers of hidden states in the best-fit models across families), mating system is not the best trait which describes these patterns. This represents a shift in understanding from previous results, in which mating system was interpreted as a clear driver of diversification rates based on elevated extinction associated with self-fertilization (Stebbins 1957; Goldberg et al. 2010).

This conclusion does not come without some limitations. This includes the described bias toward over-sampling of self-compatible systems in existing mating system data (Igić et al. 2006, Meyer et al. 2025). Additionally, the categorization of mating system as a binary value represents a simplification of continuous data. It is possible that, if further studies can incorporate quantitative data on selfing rates with appropriate SSE methods (i.e., QuaSSE; FitzJohn 2010), dynamics emerging from more granular patterns would become clear. However, regardless of the need for future study, our results question the generality of mating system as a singular driver of macroevolutionary patterns within plant reproductive biology.

### Sexual System Patterns

In contrast to mating system, sexual system more consistently affected net diversification rates. For 13 of the 16 families for which sexual system-based trait dependent models were conducted, sexual system better explained patterns of diversification compared to character-independent models. However, it is evident from our results that dioecy and monoecy do not consistently yield higher rates of diversification across families. The results from our ANOVA are not significant, pointing to the limited magnitude of many of the differences between net diversification rates. Similarly, while the resultant p-values from the chi-squared test are very near the significance threshold of 0.05, these values are still not significant.

This leads to the question: If mating system is not a strong predictor of diversification patterns, why is it that our proxy state of sexual system is a good predictor in many cases? It is possible that the binary designation of mating system as employed in this study does not appropriately capture the complex dynamics of plant mating systems. It is well-known that mating systems in plants vary considerably even within species and individual populations of the same species (Whitehead et al. 2018), and as discussed above, that the selfing-outcrossing dichotomy does not represent a true binary, but rather is a quantitative trait. To represent measures of outcrossing rate quantified by *t_m_*(the multilocus outcrossing rate) as binary data, an outcrossing rate of 0.8 is often treated as outcrossing (as in Goodwillie et al. 2005). Yet the primarily outcrossing species which this represents could still engage in some amount of selfing, despite being labeled as self-incompatible for comparative studies. This would serve as a diluting effect, since infrequent self-fertilizers could still be subject to some of the deleterious effects associated with selfing (Charlesworth and Charlesworth 1987). In contrast, while sexual systems also vary much more granularly than is captured by a three-state categorization system (reviewed in Cronk 2022), the barrier to self-fertilization presented by dioecy should be complete. This limitation is in no way unique to this study, since many studies utilize a binary framework for the assessment of mating systems. This is unsurprising, since data which captures quantitative selfing rates remains limited (Whitehead et al. 2018). This points to the importance of looking at reproductive systems holistically (in the case of this study, examining both mating and sexual systems), especially in an environment where trait information may be limited.

Another potential dynamic is that, while transitions to self-fertilization are generally irreversible (Stebbins 1957, Wright et al. 2013), transitions between sexual system states are not. For example, in Solanaceae, the reversion of dioecy back to hermaphroditism has been observed based on morphology in wild populations (Martine 2023). This has also been observed to occur rapidly through experimental evolution in Euphorbiaceae (Cossard et al. 2021). These results, and the results of our study, both suggest that dioecy may not represent an evolutionary “dead end,” as has been previously suggested as an explanation for the rarity of the reproductive system (Vamosi and Otto 2002). It is worth taking time here to evaluate this in a broader context. If we consider the hypothesis statements that self-fertilization is an evolutionary dead end (Stebbins 1957), that dioecy may have evolved to protect against the deleterious effects of self-fertilization (Charlesworth & Charlesworth 1978), and that dioecy remains uncommon due to differential extinction associated with the trait (Vamosi and Otto 2002), we are inevitably left to conclude that these three things cannot all be true in the absence of other factors.

### Conclusions and Future Directions

Reproductive traits in angiosperms are, definitionally, vital drivers of evolutionary success or failure. By understanding the macro-scale evolutionary dynamics of the past which have shaped extant angiosperm diversity, we will be better able to understand the implications of transitions to selfing and better equipped to predict the biodiversity landscape of the future.

While there have been several family-level studies on the impact of self-fertilization on diversification rates in a phylogenetic context, this represents the first study that incorporates SSE methods across replicate families in angiosperms. As noted above, the results from previous studies have been inconsistent (Marie-Orleach et al. 2024). This pattern of inconsistency persists across existing SSE studies on angiosperms (Helmstetter et al. 2022).

We draw two primary conclusions based on our results. First, we question the previous conclusion that binary mating system data can be used to explain patterns of diversification. Second, we suggest that sexual system may better explain diversification patterns than mating systems. This highlights the importance of capturing the complexity of plant reproductive biology in our analyses, and we urge future studies to carefully consider trait selection. For plant reproductive biology specifically, the integration of mating system and sexual system traits remains underutilized, despite their connected nature. Additionally, although trait data beyond broad binary classifications is often difficult to obtain, more granular observations may better capture variation for traits like mating system. Overall, these results affirm the importance of reproductive character traits for explaining patterns of diversification, which is a key goal of evolutionary biology.

## Acknowledgments

The authors would like to thank Laura F. Galloway (University of Virginia) and Rodney J. Dyer (Virginia Commonwealth University) for their support of and feedback on this project. We also thank Brian O’Meara (University of Tennessee) for several productive discussions regarding the HiSSE framework. Finally, we are grateful to the Integrative Life Sciences Doctoral Program at Virginia Commonwealth University for their ongoing support of this work.

## Author Contributions

EMM and AJE conceived of and designed the study. EMM compiled data, conducted analyses, and drafted the initial manuscript. All authors contributed to subsequent development and revisions. All authors reviewed and approved the final draft for submission.

## Data Availability

Scripts and outputs are available on GitHub at the following link: https://github.com/elenacicada/diversification_reproduction

## Conflicts of Interest

The authors have no conflicts of interest to declare.

## Funding

This work was partially supported by NSF-2423013.

---

**Table S1.**
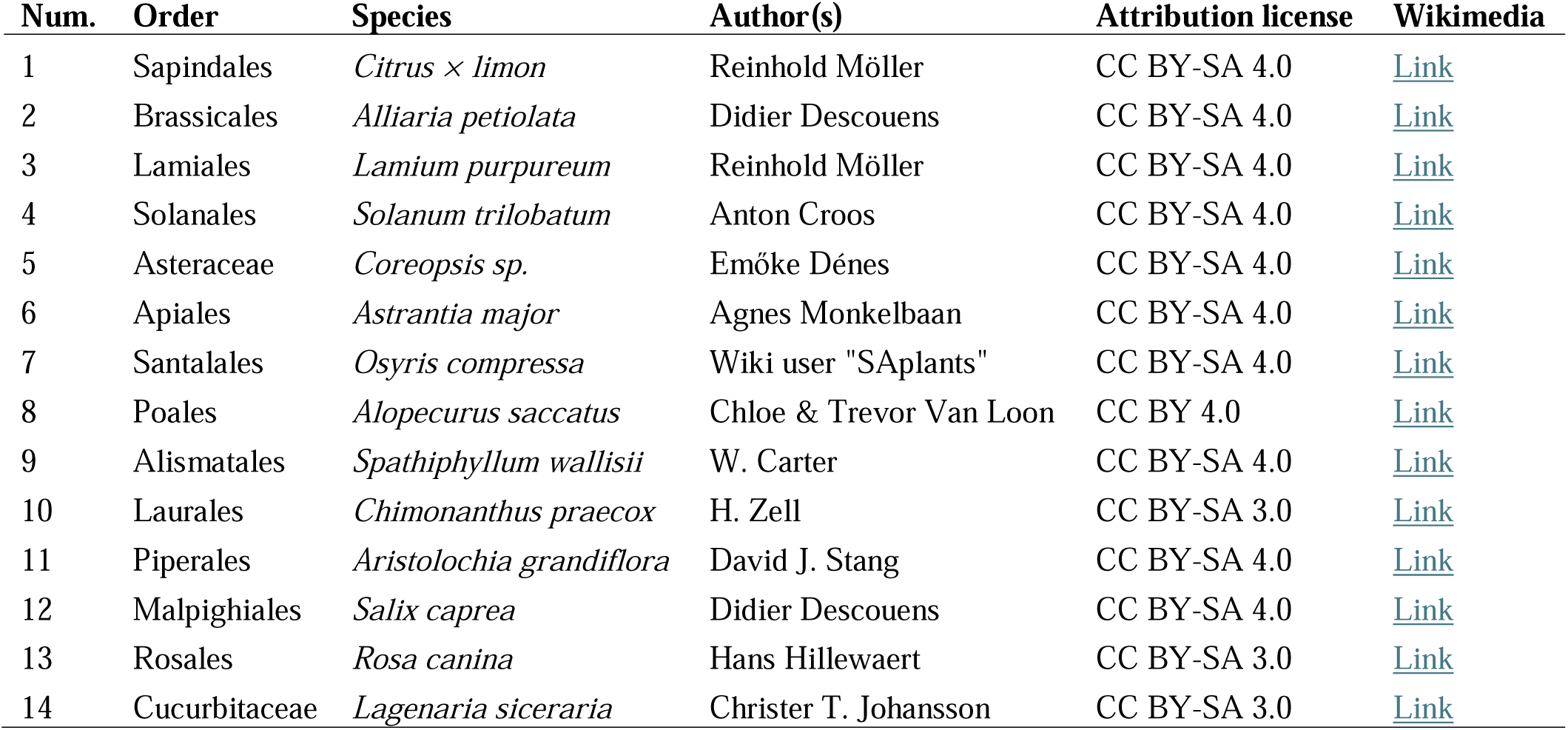
Attributions for images used in Figure 3. The first column references the number given to the taxonomic order in the figure, followed by the name of the order which the image represents, the species designation (as reported by the image author), the author’s name as listed on Wikimedia Commons, the attribution license of the work, and a direct link.

**Table S2:**
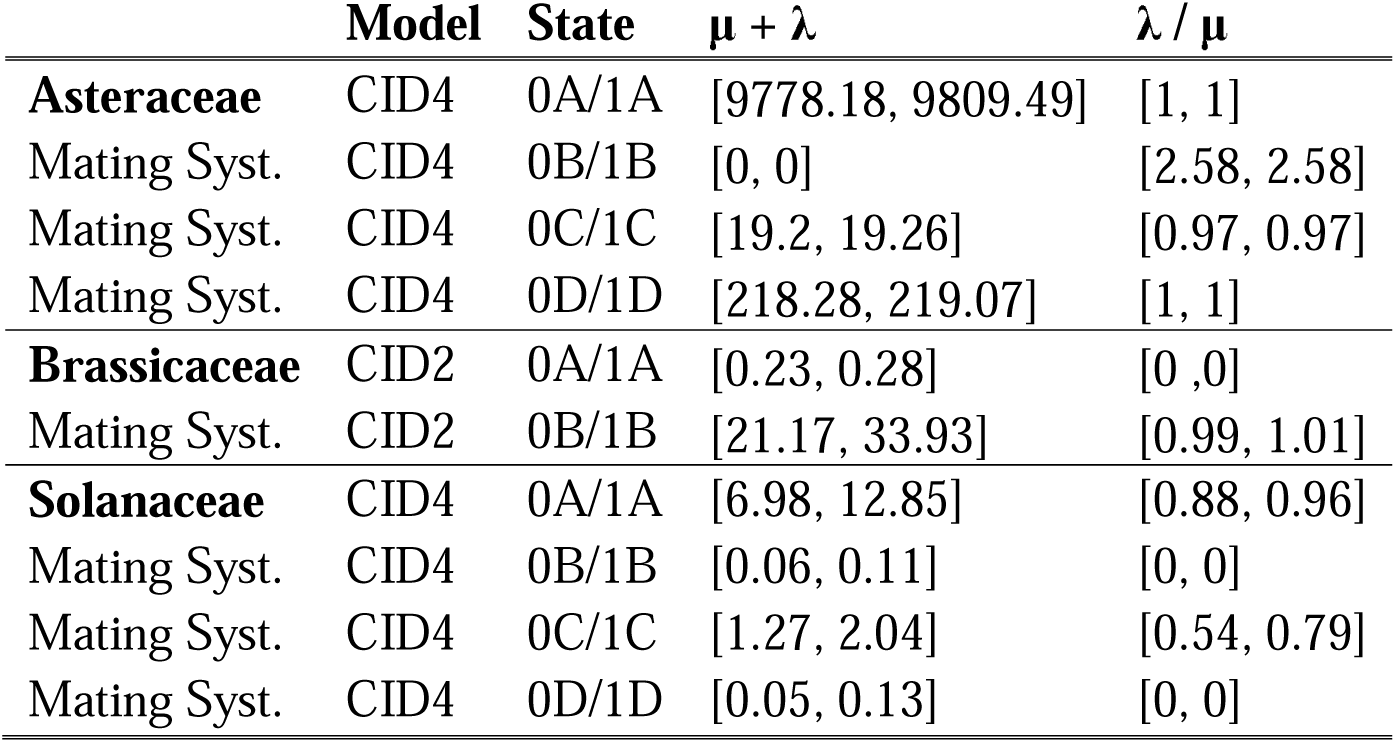
95% confidence intervals associated with mating system analysis. Letters (A-D) refer to the hidden states in the model.

**Table S3:**
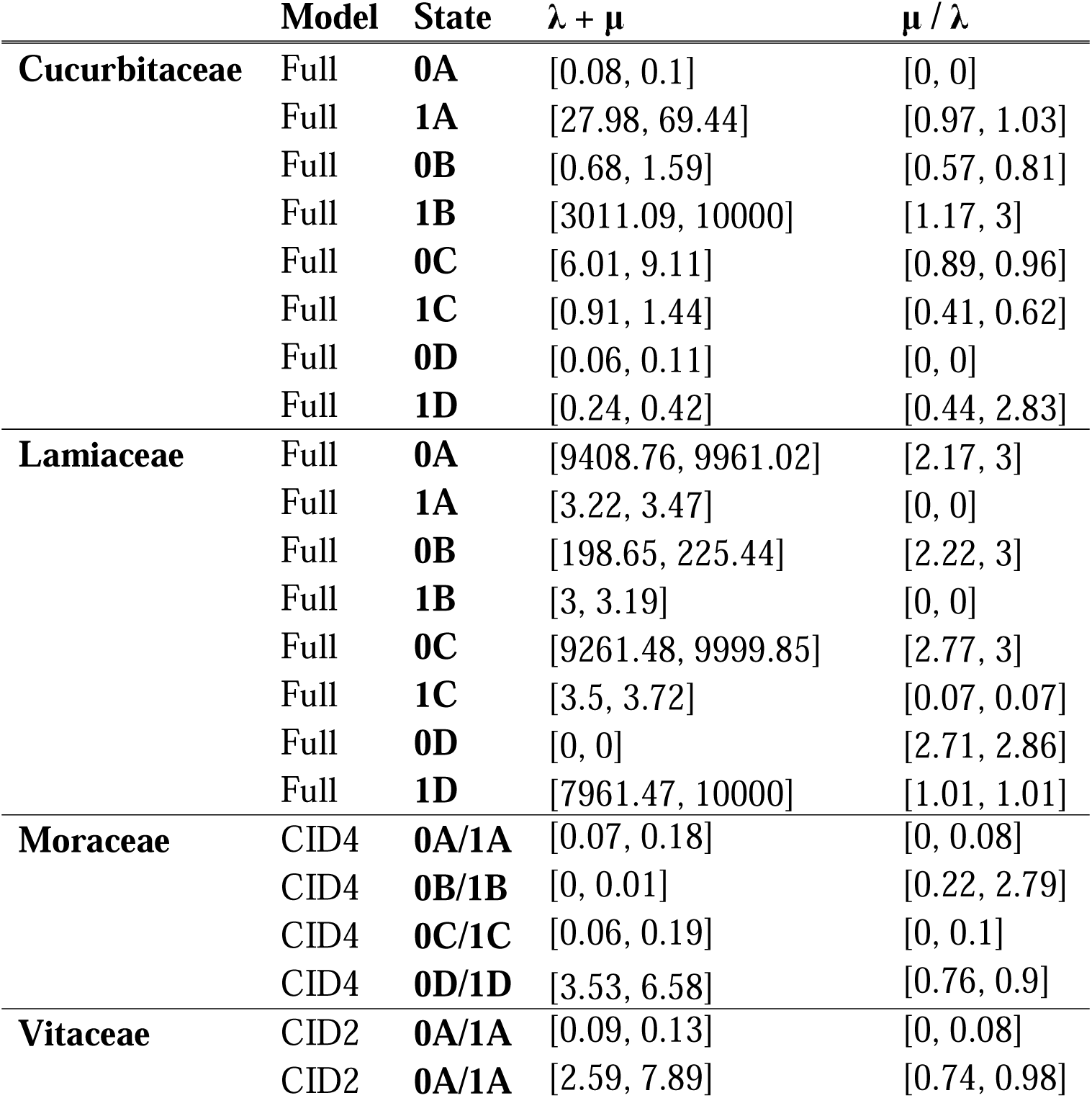
95% confidence intervals associated with two-state sexual system analysis. Letters (A-D) refer to the hidden states in the model.

**Table S4:**
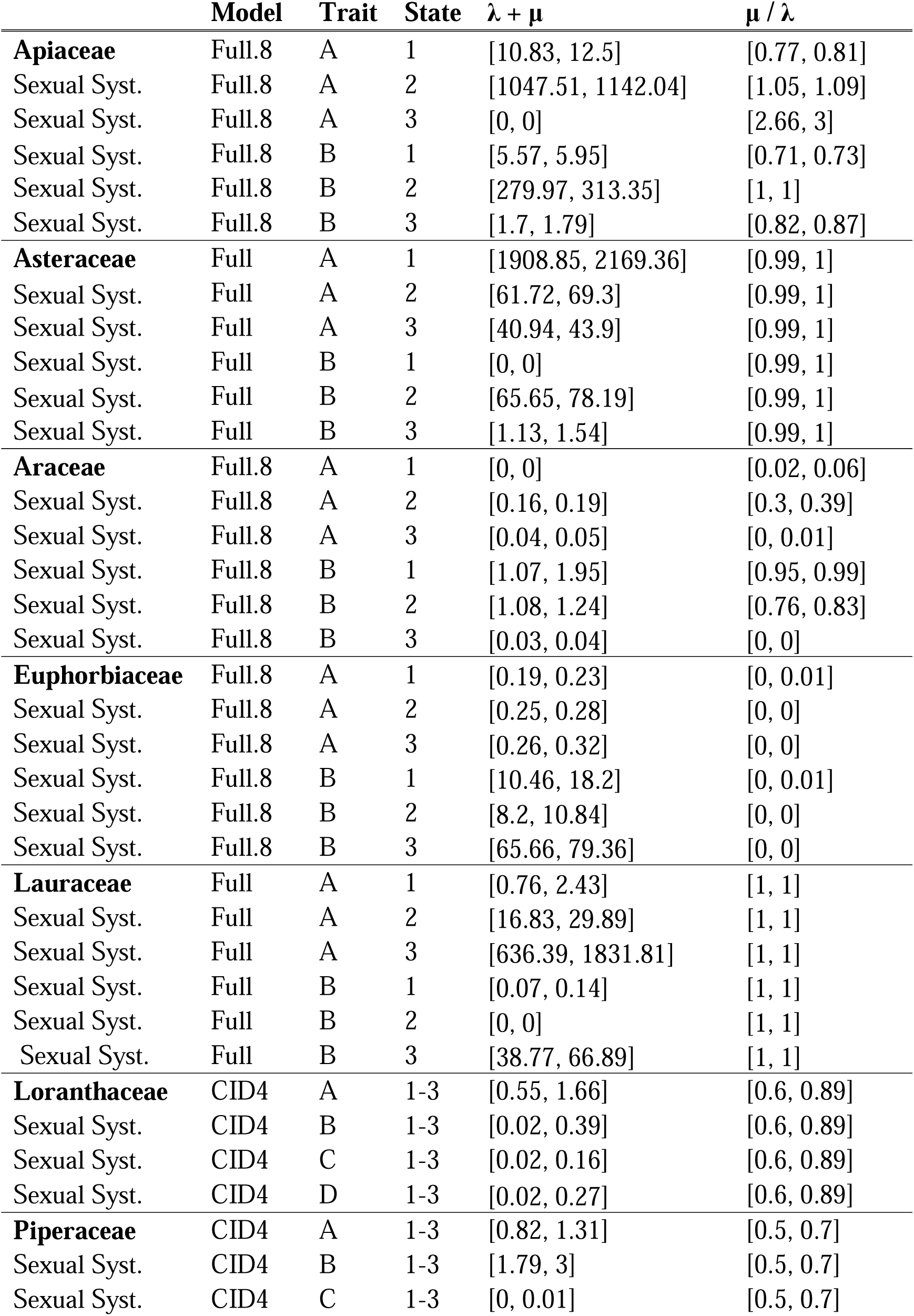
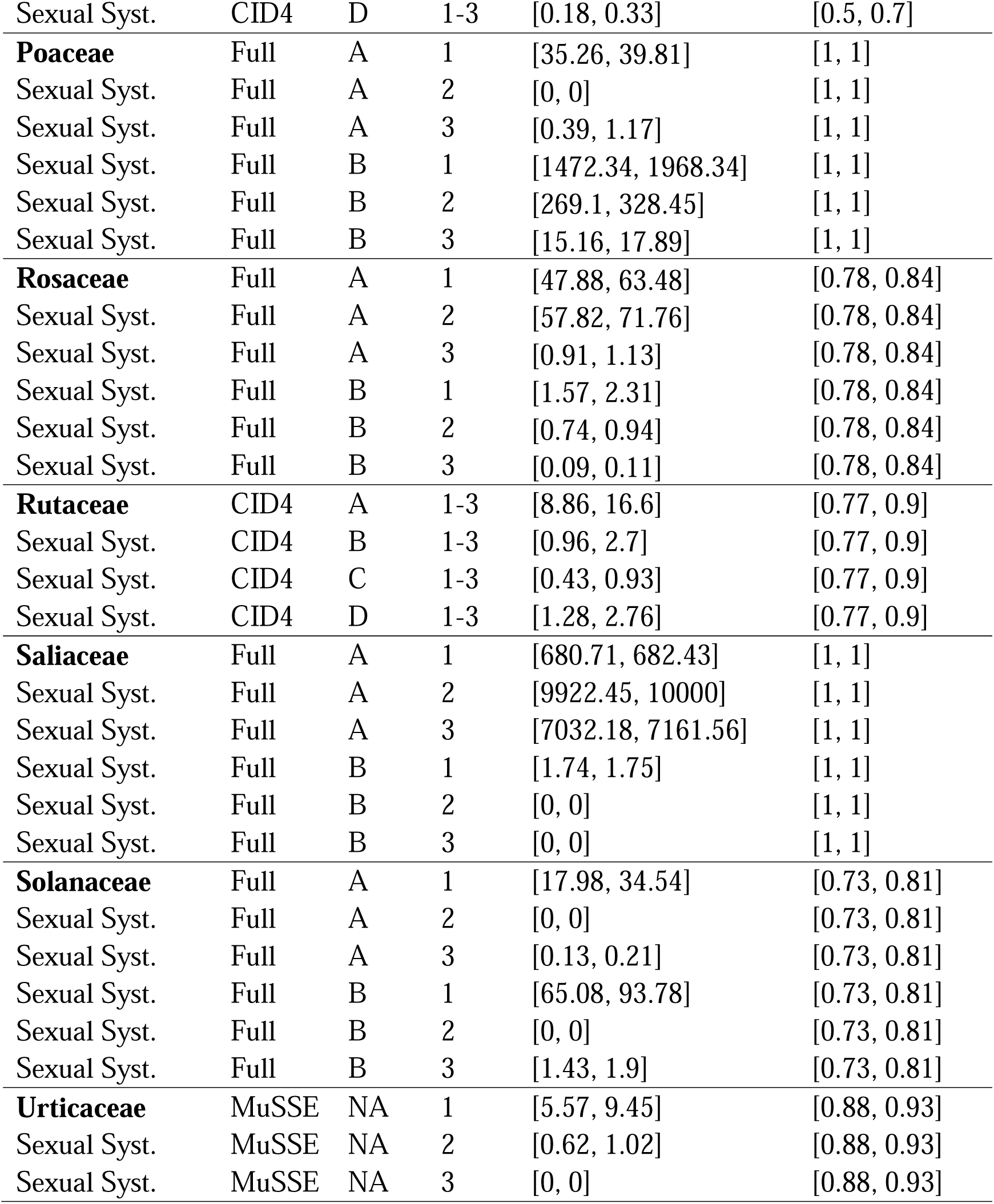
95% confidence intervals associated with sexual system analysis. Letters (A and B) refer to the hidden states in the model.

## Notes

https://github.com/elenacicada/diversification_reproduction

### Competing Interest Statement

The authors have declared no competing interest.

### Summary of Updates

Revisions were made to improve introductory materials in the paper, and to resolve minor grammar/clarity errors. No substantive changes have been made to the analysis or interpretation.

