## Supplemental Materials for "Traits and hidden states: is self-fertilization associated with rates of diversification across mating and sexual systems?"

**Table S1.** Attributions for images used in Figure 3. The first column references the number given to the taxonomic order in the figure, followed by the name of the order which the image represents, the species designation (as reported by the image author), the author's name as listed on Wikimedia Commons, the attribution license of the work, and a direct link.

| Num. | Order | Species | Author(s) | Attribution license | Wikimedia |
|---|---|---|---|---|---|
| 1 | Sapindales | *Citrus × limon* | Reinhold Möller | CC BY-SA 4.0 | Link |
| 2 | Brassicales | *Alliaria petiolata* | Didier Descouens | CC BY-SA 4.0 | Link |
| 3 | Lamiales | *Lamium purpureum* | Reinhold Möller | CC BY-SA 4.0 | Link |
| 4 | Solanales | *Solanum trilobatum* | Anton Croos | CC BY-SA 4.0 | Link |
| 5 | Asteraceae | *Coreopsis sp.* | Emőke Dénes | CC BY-SA 4.0 | Link |
| 6 | Apiales | *Astrantia major* | Agnes Monkelbaan | CC BY-SA 4.0 | Link |
| 7 | Santalales | *Osyris compressa* | Wiki user "SAplants" | CC BY-SA 4.0 | Link |
| 8 | Poales | *Alopecurus saccatus* | Chloe & Trevor Van Loon | CC BY 4.0 | Link |
| 9 | Alismatales | *Spathiphyllum wallisii* | W. Carter | CC BY-SA 4.0 | Link |
| 10 | Laurales | *Chimonanthus praecox* | H. Zell | CC BY-SA 3.0 | Link |
| 11 | Piperales | *Aristolochia grandiflora* | David J. Stang | CC BY-SA 4.0 | Link |
| 12 | Malpighiales | *Salix caprea* | Didier Descouens | CC BY-SA 4.0 | Link |
| 13 | Rosales | *Rosa canina* | Hans Hillewaert | CC BY-SA 3.0 | Link |
| 14 | Cucurbitaceae | *Lagenaria siceraria* | Christer T. Johansson | CC BY-SA 3.0 | Link |

**Table S2:** 95% confidence intervals associated with mating system analysis.

| | **Model** | **State** | **μ + λ** | **λ / μ** |
|---|---|---|---|---|
| **Asteraceae** | CID4 | 0A/1A | [9778.18, 9809.49] | [1, 1] |
| Mating Syst. | CID4 | 0B/1B | [0, 0] | [2.58, 2.58] |
| Mating Syst. | CID4 | 0C/1C | [19.2, 19.26] | [0.97, 0.97] |
| Mating Syst. | CID4 | 0D/1D | [218.28, 219.07] | [1, 1] |
| **Brassicaceae** | CID2 | 0A/1A | [0.23, 0.28] | [0 ,0] |
| Mating Syst. | CID2 | 0B/1B | [21.17, 33.93] | [0.99, 1.01] |
| **Solanaceae** | CID4 | 0A/1A | [6.98, 12.85] | [0.88, 0.96] |
| Mating Syst. | CID4 | 0B/1B | [0.06, 0.11] | [0, 0] |
| Mating Syst. | CID4 | 0C/1C | [1.27, 2.04] | [0.54, 0.79] |
| Mating Syst. | CID4 | 0D/1D | [0.05, 0.13] | [0, 0] |

**Table S3:** 95% confidence intervals associated with two-state sexual system analysis.

| | Model | State | λ + μ | μ / λ |
|---|---|---|---|---|
| **Cucurbitaceae** | Full | **0A** | [0.08, 0.1] | [0, 0] |
| | Full | **1A** | [27.98, 69.44] | [0.97, 1.03] |
| | Full | **0B** | [0.68, 1.59] | [0.57, 0.81] |
| | Full | **1B** | [3011.09, 10000] | [1.17, 3] |
| | Full | **0C** | [6.01, 9.11] | [0.89, 0.96] |
| | Full | **1C** | [0.91, 1.44] | [0.41, 0.62] |
| | Full | **0D** | [0.06, 0.11] | [0, 0] |
| | Full | **1D** | [0.24, 0.42] | [0.44, 2.83] |
| **Lamiaceae** | Full | **0A** | [9408.76, 9961.02] | [2.17, 3] |
| | Full | **1A** | [3.22, 3.47] | [0, 0] |
| | Full | **0B** | [198.65, 225.44] | [2.22, 3] |
| | Full | **1B** | [3, 3.19] | [0, 0] |
| | Full | **0C** | [9261.48, 9999.85] | [2.77, 3] |
| | Full | **1C** | [3.5, 3.72] | [0.07, 0.07] |
| | Full | **0D** | [0, 0] | [2.71, 2.86] |
| | Full | **1D** | [7961.47, 10000] | [1.01, 1.01] |
| **Moraceae** | CID4 | **0A/1A** | [0.07, 0.18] | [0, 0.08] |
| | CID4 | **0B/1B** | [0, 0.01] | [0.22, 2.79] |
| | CID4 | **0C/1C** | [0.06, 0.19] | [0, 0.1] |
| | CID4 | **0D/1D** | [3.53, 6.58] | [0.76, 0.9] |
| **Vitaceae** | CID2 | **0A/1A** | [0.09, 0.13] | [0, 0.08] |
| | CID2 | **0A/1A** | [2.59, 7.89] | [0.74, 0.98] |

**Table S4:** 95% confidence intervals associated with sexual system analysis.

| | **Model** | **Trait** | **State** | **λ + μ** | **μ / λ** |
|---|---|---|---|---|---|
| **Apiaceae** | Full.8 | A | 1 | [10.83, 12.5] | [0.77, 0.81] |
| Sexual Syst. | Full.8 | A | 2 | [1047.51, 1142.04] | [1.05, 1.09] |
| Sexual Syst. | Full.8 | A | 3 | [0, 0] | [2.66, 3] |
| Sexual Syst. | Full.8 | B | 1 | [5.57, 5.95] | [0.71, 0.73] |
| Sexual Syst. | Full.8 | B | 2 | [279.97, 313.35] | [1, 1] |
| Sexual Syst. | Full.8 | B | 3 | [1.7, 1.79] | [0.82, 0.87] |
| **Asteraceae** | Full | A | 1 | [1908.85, 2169.36] | [0.99, 1] |
| Sexual Syst. | Full | A | 2 | [61.72, 69.3] | [0.99, 1] |
| Sexual Syst. | Full | A | 3 | [40.94, 43.9] | [0.99, 1] |
| Sexual Syst. | Full | B | 1 | [0, 0] | [0.99, 1] |
| Sexual Syst. | Full | B | 2 | [65.65, 78.19] | [0.99, 1] |
| Sexual Syst. | Full | B | 3 | [1.13, 1.54] | [0.99, 1] |
| **Araceae** | Full.8 | A | 1 | [0, 0] | [0.02, 0.06] |
| Sexual Syst. | Full.8 | A | 2 | [0.16, 0.19] | [0.3, 0.39] |
| Sexual Syst. | Full.8 | A | 3 | [0.04, 0.05] | [0, 0.01] |
| Sexual Syst. | Full.8 | B | 1 | [1.07, 1.95] | [0.95, 0.99] |
| Sexual Syst. | Full.8 | B | 2 | [1.08, 1.24] | [0.76, 0.83] |
| Sexual Syst. | Full.8 | B | 3 | [0.03, 0.04] | [0, 0] |
| **Euphorbiaceae** | Full.8 | A | 1 | [0.19, 0.23] | [0, 0.01] |
| Sexual Syst. | Full.8 | A | 2 | [0.25, 0.28] | [0, 0] |
| Sexual Syst. | Full.8 | A | 3 | [0.26, 0.32] | [0, 0] |
| Sexual Syst. | Full.8 | B | 1 | [10.46, 18.2] | [0, 0.01] |
| Sexual Syst. | Full.8 | B | 2 | [8.2, 10.84] | [0, 0] |
| Sexual Syst. | Full.8 | B | 3 | [65.66, 79.36] | [0, 0] |
| **Lauraceae** | Full | A | 1 | [0.76, 2.43] | [1, 1] |
| Sexual Syst. | Full | A | 2 | [16.83, 29.89] | [1, 1] |
| Sexual Syst. | Full | A | 3 | [636.39, 1831.81] | [1, 1] |
| Sexual Syst. | Full | B | 1 | [0.07, 0.14] | [1, 1] |
| Sexual Syst. | Full | B | 2 | [0, 0] | [1, 1] |
| Sexual Syst. | Full | B | 3 | [38.77, 66.89] | [1, 1] |
| **Loranthaceae** | CID4 | A | 1-3 | [0.55, 1.66] | [0.6, 0.89] |
| Sexual Syst. | CID4 | B | 1-3 | [0.02, 0.39] | [0.6, 0.89] |
| Sexual Syst. | CID4 | C | 1-3 | [0.02, 0.16] | [0.6, 0.89] |
| Sexual Syst. | CID4 | D | 1-3 | [0.02, 0.27] | [0.6, 0.89] |
| **Piperaceae** | CID4 | A | 1-3 | [0.82, 1.31] | [0.5, 0.7] |
| Sexual Syst. | CID4 | B | 1-3 | [1.79, 3] | [0.5, 0.7] |
| Sexual Syst. | CID4 | C | 1-3 | [0, 0.01] | [0.5, 0.7] |
| Sexual Syst. | CID4 | D | 1-3 | [0.18, 0.33] | [0.5, 0.7] |

| | | | | | |
|---|---|---|---|---|---|
| **Poaceae** | Full | A | 1 | [35.26, 39.81] | [1, 1] |
| Sexual Syst. | Full | A | 2 | [0, 0] | [1, 1] |
| Sexual Syst. | Full | A | 3 | [0.39, 1.17] | [1, 1] |
| Sexual Syst. | Full | B | 1 | [1472.34, 1968.34] | [1, 1] |
| Sexual Syst. | Full | B | 2 | [269.1, 328.45] | [1, 1] |
| Sexual Syst. | Full | B | 3 | [15.16, 17.89] | [1, 1] |
| **Rosaceae** | Full | A | 1 | [47.88, 63.48] | [0.78, 0.84] |
| Sexual Syst. | Full | A | 2 | [57.82, 71.76] | [0.78, 0.84] |
| Sexual Syst. | Full | A | 3 | [0.91, 1.13] | [0.78, 0.84] |
| Sexual Syst. | Full | B | 1 | [1.57, 2.31] | [0.78, 0.84] |
| Sexual Syst. | Full | B | 2 | [0.74, 0.94] | [0.78, 0.84] |
| Sexual Syst. | Full | B | 3 | [0.09, 0.11] | [0.78, 0.84] |
| **Rutaceae** | CID4 | A | 1-3 | [8.86, 16.6] | [0.77, 0.9] |
| Sexual Syst. | CID4 | B | 1-3 | [0.96, 2.7] | [0.77, 0.9] |
| Sexual Syst. | CID4 | C | 1-3 | [0.43, 0.93] | [0.77, 0.9] |
| Sexual Syst. | CID4 | D | 1-3 | [1.28, 2.76] | [0.77, 0.9] |
| **Saliaceae** | Full | A | 1 | [680.71, 682.43] | [1, 1] |
| Sexual Syst. | Full | A | 2 | [9922.45, 10000] | [1, 1] |
| Sexual Syst. | Full | A | 3 | [7032.18, 7161.56] | [1, 1] |
| Sexual Syst. | Full | B | 1 | [1.74, 1.75] | [1, 1] |
| Sexual Syst. | Full | B | 2 | [0, 0] | [1, 1] |
| Sexual Syst. | Full | B | 3 | [0, 0] | [1, 1] |
| **Solanaceae** | Full | A | 1 | [17.98, 34.54] | [0.73, 0.81] |
| Sexual Syst. | Full | A | 2 | [0, 0] | [0.73, 0.81] |
| Sexual Syst. | Full | A | 3 | [0.13, 0.21] | [0.73, 0.81] |
| Sexual Syst. | Full | B | 1 | [65.08, 93.78] | [0.73, 0.81] |
| Sexual Syst. | Full | B | 2 | [0, 0] | [0.73, 0.81] |
| Sexual Syst. | Full | B | 3 | [1.43, 1.9] | [0.73, 0.81] |
| **Urticaceae** | MuSSE | NA | 1 | [5.57, 9.45] | [0.88, 0.93] |
| Sexual Syst. | MuSSE | NA | 2 | [0.62, 1.02] | [0.88, 0.93] |
| Sexual Syst. | MuSSE | NA | 3 | [0, 0] | [0.88, 0.93] |
